# Prefrontal pathways provide top down control of memory for sequences of events

**DOI:** 10.1101/508051

**Authors:** Maanasa Jayachandran, Stephanie Linley, Maximilian Schlecht, Stephen V. Mahler, Robert P. Vertes, Timothy A. Allen

**Affiliations:** Cognitive Neuroscience Program, Department of Psychology, Florida International University, Miami, FL, 33199, USA; Center for Complex Systems and Brain Sciences, Florida Atlantic University, Boca Raton, FL, 33431, USA; Neurobiology and Behavior, University of California Irvine, Irvine, CA, 92697, USA; Enviromental Health Sciences, Robert Stempel College of Public Health, Florida International University, Miami, FL, 33199, USA

**Keywords:** sequence memory, temporal context, episodic memory, DREADDs, back projections, nucleus reuniens, perirhinal cortex

## Abstract

We remember our lives as sequences of events, but it is unclear how these memories are controlled during retrieval. In rats, prelimbic cortex (PL) is positioned to influence sequence memory through extensive top down inputs to the nucleus reuniens of the thalamus (RE) and perirhinal cortex (PER), regions heavily interconnected with the hippocampus. Here, we tested the hypothesis that specific PL➔RE and PL➔PER projections regulate sequence memory retrieval using an hM4Di synaptic-silencing approach. First, we show that the suppression of PL activity impairs sequence memory. Second, we show that inhibiting PL➔RE and PL➔PER pathways effectively eliminated sequence memory. Last, we performed a sequential lag analysis showing that the PL➔RE pathway contributes to a working memory retrieval strategy, and the PL➔PER pathway contributes to a temporal context memory retrieval strategy. These results demonstrate that the PL➔RE and PL➔PER pathways serve as top down mechanisms that control sequence memory retrieval strategies.

## Introduction

We remember our lives as sequences of events, an ability at the core of episodic memory. The temporal organization of memory has been studied in a wide array of tasks and is generally thought to be useful for disambiguating memories with overlapping content (Clayton and Dickinson, 1998; Henson, 2001; Tulving, 2002; Agster et al., 2002; Allen and Fortin, 2013; Eichenbaum, 2017a).

Neurobiologically, memory for sequences of events relies on the hippocampus (Fortin et al., 2002; Kesner et al., 2002; Allen et al., 2016) and medial prefrontal cortex (Barker et al., 2007; Euston et al., 2007; Hales et al., 2009; Blumenfeld et al., 2011; Tiganj et al., 2018). However, it is clear that these regions serve different roles in sequence memory (Hsieh and Ranganath, 2015; Reeders et al., 2018). The hippocampus is thought to associate information with spatiotemporal contexts (Eichenbaum, 2004; Knierim et al., 2006; Knierim, 2015; Skelin et al., 2018), whereas PL is thought to influence the rapid retrieval of information relevant to an action or decision (Ferbinteanu et al., 2006; Euston et al., 2012; Preston and Eichenbaum, 2013).

In rats, PL is ideally situated to influence memory retrieval through its many projections to the thalamus and cortex (Sesack et al., 1989; Chiba et al., 2001; Vertes, 2002; Hoover and Vertes, 2007). One idea is that PL can select or bias different sequence retrieval strategies through its top down projection pathways. If true, we should be able to impair sequence memory through the selective inhibition of different PL projection pathways with somewhat different effects in each pathway.

In sequence memory, the most significant PL pathways target the nucleus reuniens of the thalamus (RE) and perirhinal cortex (PER) because they are strongly interconnected with the hippocampus (Eichenbaum, 2017b). RE receives diverse and widely distributed projections from limbic-related sites (Vertes, 2002; 2004; McKenna and Vertes, 2004), however, RE projections are relatively restricted and primarily bound for the hippocampus, parahippocampus, and prefrontal cortex (Vertes, 2006; Vertes et al., 2006). Given the absence of direct medial prefrontal cortex to hippocampus projections, RE represents the primary route as follows: HC➔mPFC➔RE➔HC (Vertes et al., 2007). Notably, RE is critical to a variety of working memory tasks that have been linked with medial prefrontal cortex-associated deficits (Davoodi et al., 2009; Dolleman-van der weel et al., 2009; Cassel et al., 2013; Hallock et al., 2013; Xu and Südhof, 2013; Griffin, 2015; Ito et al., 2015; Viena et al., 2018). Like RE, PER sits at a nexus of bidirectional communication between the hippocampus and PL (Furtak et al., 2007). While PER is known to be critical to memory for complex stimuli (Murray et al., 2000; Bussey et al., 2002; Aggleton et al., 2010; Barker and Warburton, 2011; Feinberg et al., 2012), PER also appears to be critical for temporal aspects of memory (Murray and Richmond, 2001; Bussey et al., 2005; Allen et al., 2007; Barker et al., 2007; Bang and Brown, 2009; Chen et al., 2015; Naya et al., 2017).

Here we tested the role of these two PL projection pathways using a novel sequence memory task that taxes memory for the order of events and allows an analysis of the underlying strategy (Allen et al., 2014; 2016). Briefly, rats sampled two sequences of odors and demonstrated sequence memory by indicating whether each odor was presented ‘in sequence’ (InSeq) or ‘out of sequence’ (OutSeq). First, we tested the role of PL in sequence memory using a chemogenetic approach via systemic intraperitoneal (i.p.) injections of clozapine-n-oxide (CNO) in rats expressing inhibitory DREADDs (designer receptor exclusively activated by designer drugs; hM4Di). Second, we tested the role of PL➔RE and PL➔PER pathways using a synaptic silencing approach (Mahler et al., 2014; Stachniak et al., 2014; Roth, 2016; Smith et al., 2016; Lichtenberg et al., 2017). Last, we tested the unique contributions of PL➔RE and PL➔ PER pathways using a lag analysis across specific probes (e.g., AB<u>A</u>, where A is a repeated item from two positions earlier, or AB<u>D</u>, where D skips ahead one position). Theoretically, different lag performance patterns on repeated items can distinguish contributions of working memory and temporal context memory (see Reeders et al., 2018).

The results show that PL is critical to sequence memory, that silencing PL➔RE and PL➔PER projections effectively eliminates sequence memory, and that each projection leads to a unique pattern of behavioral deficits. Silencing PL➔RE pathways produced a reduction in working memory, whereas, silencing PL➔PER pathways produced a reduction in temporal context memory (i.e., graded memory retrieval based on the temporal proximity of items in a sequence). These results advance the concept that PL inputs to RE and PER provide top down control of sequence memory, and can regulate memory retrieval strategies.

## Results

### Sequence memory

We trained rats on a modified version of an odor sequence task (Allen et al., 2016). The behavioral apparatus was fully automated and capable of repeated deliveries of multiple distinct odors (pure chemical odorants) between two odor ports. Here we used two sequences, each composed of four odors that occurred at the opposite ends of a straight alley maze (Figure 1A). By using two sequences (compared to only one sequence in previous task versions) we eliminated the possibility that the rats can hold a single sequence in memory throughout testing, as well as increased the overall memory load. This allowed us to tease apart the use of different retrieval strategies as rats had to retrieve sequences repeatedly from long term memory stores. Rats demonstrated sequence memory by indicating whether odors were InSeq (holding in nose port >1s) or OutSeq (withdrawing from nose port <1s) within a sequence in order to receive a water reward (Figure 1B and 1C). We calculated a sequence memory index (SMI), which measures overall sequence memory while controlling for individual differences in poking behavior (Allen et al., 2014). The SMI normalizes the proportion of InSeq and OutSeq items presented during a single session for comparison across sessions, and scores sequence memory with a value ranging from −1 to 1. A score of 0 indicates chance performance. A score of 1 is perfect performance. Rats were trained on the sequence memory task over several weeks in progressive stages as depicted in Figure 1F.

**Figure 1.**
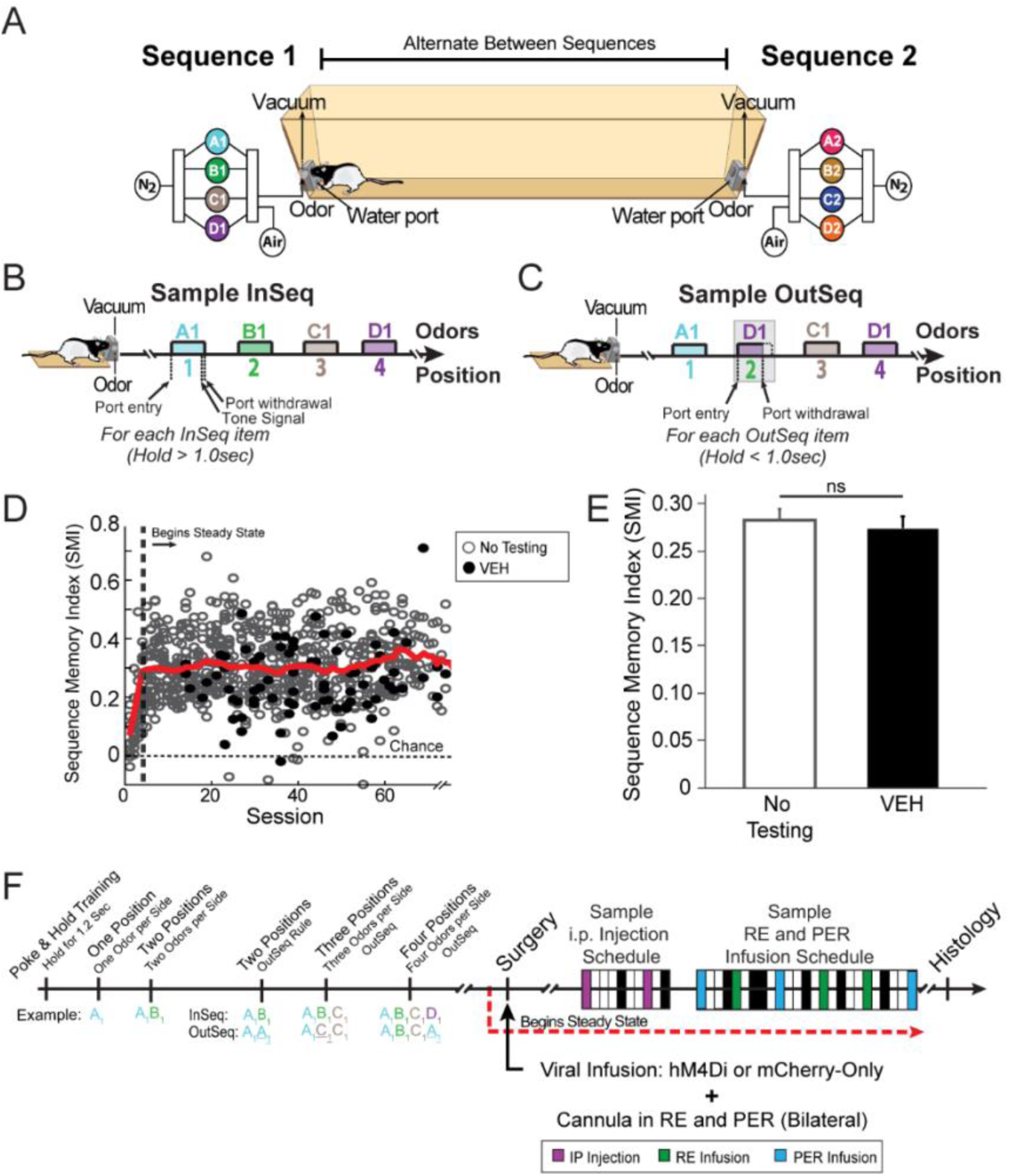
Sequence memory task. The sequence memory task tests the ability to remember the order of events within a sequence. (A) The apparatus comprises of a linear track with odor ports at each end where rats were presented with two separate four-odor sequences (A_1_, B_1_, C_1_, D_1_ or A_2_, B_2_, C_2_, D_2_). In a session, the two sequences were alternated with up to 100 sequence presentations per session. The use of two sequences (compared to previous versions using a single sequence; Allen et al., 2014; Allen et al., 2016) reduces the possibility that the rats keep one sequence in mind throughout the session and adds to the overall memory load. (B and C) Individual odor presentations were initiated by a nose poke, and rats were required to correctly identify the odor as either InSeq (by holding >1s) or OutSeq (by holding <1s) to receive a water reward. (B) Each session consisted of approximately 70% of the odors being InSeq (e.g., ABCD) and (C) 30% of one odor within a sequence being OutSeq (e.g, A**<u>D</u>**CD). (D) We used a sequence memory index (SMI; see methods) to collapse the behavioral data of each session into a single normalized measure of sequence memory performance. An SMI value of 1 represents perfect performance (correctly detecting InSeq and OutSeq items), while 0 represents chance level performance. The open gray circles represent the no testing sessions and closed black circles represent the vehicle sessions. The red line represents the mean SMI of both no testing and vehicle combined (calculated with a sliding window of 10 sessions) as the rats learned the task. Steady state was reached approximately five sessions after the rats were introduced to both of the full sequences. (E) There was no significant difference between no testing and vehicle sessions. (F) Rats were trained on both sequences until they reached steady state performance levels. We focused our analysis on two experimental blocks, i) i.p. injection suppressing PL neurons (magenta), and ii) cannula infusions targeting PL terminals in RE (green) or PER (blue). The boxes represent a sample schedule of both vehicle and CNO days. Vehicle days are denoted in black and no testing days are denoted in white. The order was randomized and counterbalanced between rats. Abbreviations: InSeq, in sequence item(s); OutSeq, out of sequence item(s); PER, perirhinal cortex; RE, nucleus reuniens of the thalamus; VEH, vehicle; i.p., intraperitoneal; SMI, sequence memory index; ns = not significant.

Overall, rats demonstrated strong sequence memory (SMI_well-trained_: 0.302 ± 0.034) with no significant differences between no testing days and vehicle days (Figure 1E; t_(90)_ = 0.426, p = 0.673). Expected vs. observed frequencies were analyzed with G-tests to determine whether the observed frequency of InSeq and OutSeq responses for a given session was significantly different than chance. Single-subject analyses showed that every rat differentiated InSeq and OutSeq items at levels well above chance (Figure S2; all G-tests had p’s < 0.05). Moreover, rats performed well on position 2 (SMI_Pos2_: 0.267 ± 0.021), position 3 (SMI_Pos3_: 0.295 ± 0.023), and position 4 (SMI_Pos4_: 0.213 ± 0.042) indicating memory for the entire length of each sequence. Position1 was excluded since an OutSeq item could never be presented in that position. Rats also performed well above chance levels on sequence1 (SMI_Seq1_: 0.285 ± 0.009) and sequence2 (SMI_Seq2_: 0.270 ± 0.009), with no significant difference between sequences (t_(704)_ = 1.412, p = 0.158), indicating that rats were successfully switching between the two sequences.

Additionally, performance was not significantly different between no testing days and vehicle days on either of the sequences (Figure S3A; sequence1: t_(12)_ = 0.209, p = 0.838; sequence2: t_(12)_ = 0.566, p = 0.586).

### hM4Di in PL neurons

We targeted PL, the most implicated subregion of the rodent medial prefrontal cortex for the temporal organization of memory (Uylings et al., 2003; DeVito and Eichenbaum, 2011; Tiganj et al., 2017; 2018). After reaching behavioral criterion (asymptotic sequence memory performance levels over multiple sessions) rats underwent surgery for microinjection of one of two viral construct groups (hM4Di+: AAV9.CAG.mCherry-2a-hM4D _i_^nrxn^.WPRE.SV40; or mCherry-only: AAV9.CB7.CI.mCherry.WPRE.rBG) into PL bilaterally (A/P: 3.24mm, M/L: ± 0.7mm, D/V_from cortex_: −2.8mm). A modified hM4Di was used, which is as an axon-preferring variant referred to as hM4D_i_^nrxn^ (neurexin; Stachniak et al., 2014). This particular variant has been shown to exhibit enhanced axonal expression and reduced somatic expression. The activation of this hM4D_i_^nrxn^ variant inhibits synaptic transmission without somatic hyperpolarization (Stachniak et al., 2014). In addition to the viral infusion, chronic cannula were implanted targeting RE (at a 10° angle in order to avoid the superior sagittal sinus; A/P −1.8 mm, M/L −1.2 mm, D/V −6.7 mm) and PER bilaterally (A/P −6.0 mm, M/L ±6.8 mm, D/V −6.0 mm). The postsurgical viral gestation time for these experiments was determined by injecting hM4Di in a non-behavioral group of rats (n = 4) perfused at a series of time points (1, 2, 4, and 8 weeks) following surgery. At two weeks the virus was well expressed (Figure S1), thus our experiments began after this incubation period. After recovery, rats continued performing the sequence task daily and the experimental manipulations (i.p. injections and cannula infusions) began ∼3 weeks post cannulation/viral injection.

### Localization of AAV infection and hM4Di expression

We visualized the expression resulting from both AAV viral constructs immunohistochemically using antisera for the fluorescent reporter mCherry. The spread of hM4Di+ neurons across the prefrontal cortex was precisely mapped onto a series of schematic plates for all rats, shown in Figure 3B and S4. hM4Di+ cells were concentrated in the rostral to mid-levels of PL and the anterior cingulate cortex, with dense labeling seen in deep cortical layers. While mCherry labeled cells were present throughout the rostral caudal extent of PL, hM4Di+ expression in anterior cingulate cortex was limited to rostral aspects, with only a scarce number of infected cells present in the posterior and ventral divisions of the anterior cingulate cortex. In some hM4Di+ cases, there were negligible numbers of labeled cells which extended into the medial orbital, ventral orbital, infralimbic, medial agranular cortex, and claustrum.

To estimate the localization of viral transduction with hM4Di+, we processed a series of slices for dual immunofluorescence in a subset of animals (n = 3) using antisera for mCherry and NeuN (Figure 2A). Cell counts were conducted in FIJI for mCherry labeled neurons and NeuN labeled neurons at three anterior posterior (AP) levels: at the core of the injection site of the viral vector in layers 5/6 of PL and anterior cingulate cortex (AP +3.2); and anterior and posterior to this level (± 120um), allowing for a ratio of mCherry to NeuN labeled cells to determine the percentage of hM4Di+ infected cells. Figure 2B depicts the mean percent of double-labeled cells in PL and anterior cingulate cortex at each of these levels. Overall, the maximum percentage of hM4Di+ infected neurons in PL (48.99% ± 4.32) and anterior cingulate cortex (50.61% ± 3.93) did not differ across subdivisions (t_(16)_ = 0.779, p = 0.785) such that approximately ∼50% of neurons were double labeled.

**Figure 2.**
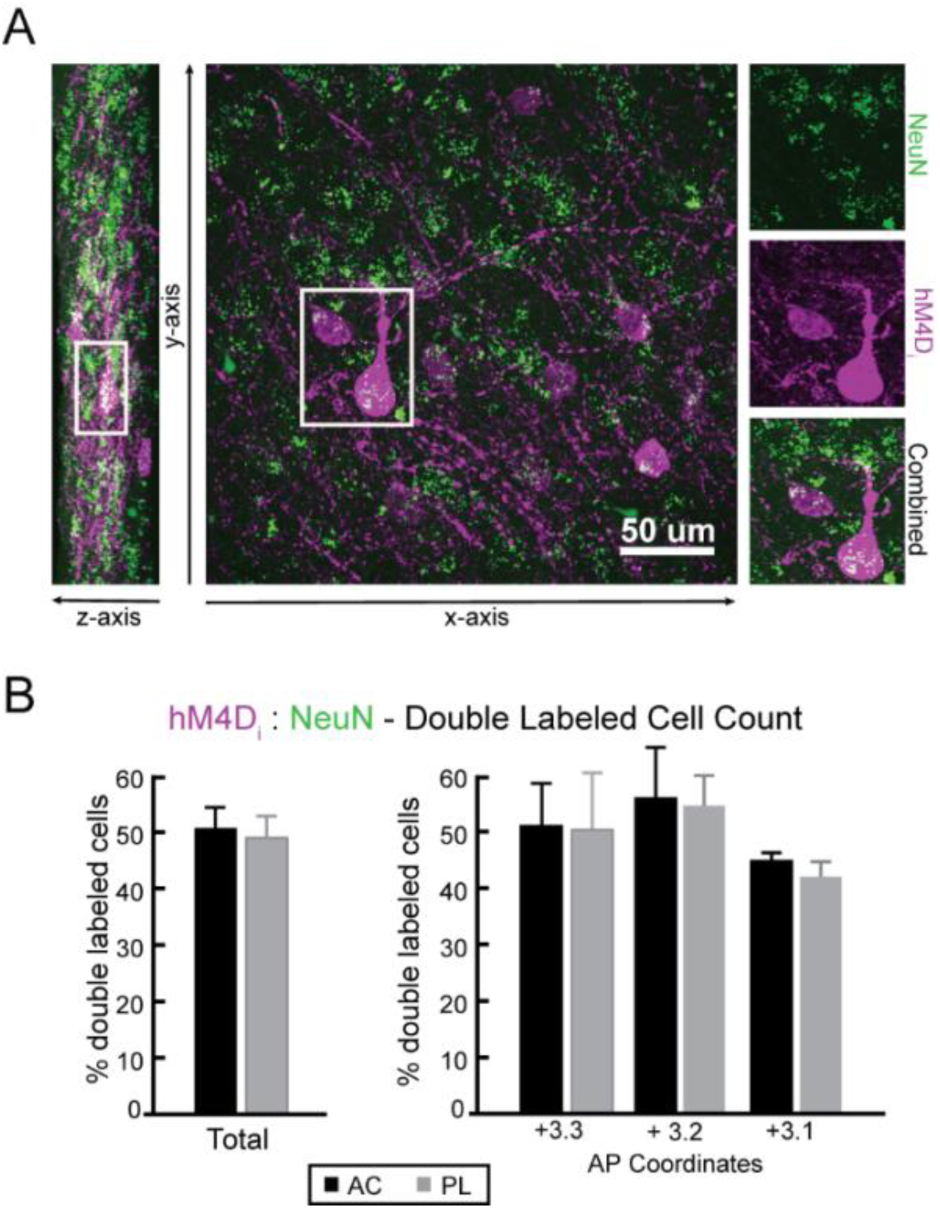
hM4Di expression in prelimbic cortex. The quantification of h4MDi infected neurons in PL. (A) A confocal image demonstrates how double labeled mCherry and NeuN (neuron-specific) cells were quantified. Boxed area represents a double-labeled cell. (B) h4MDi infection rates in PL and anterior cingulate cortex were determined by performing cell counts of mCherry and NeuN double labeled neurons for a subset of rats (n=3) at three anterior posterior levels. The mean percent of double labeled cells (± SEM) was calculated by dividing the total number of mCherry/NeuN labeled cells divided by the total number of NeuN cells across a uniform region of interest. Abbreviations: PL, prelimbic cortex; AC, anterior cingulate cortex; NeuN, neuronal nuclei; AP, anterior posterior.

**Figure 3.**
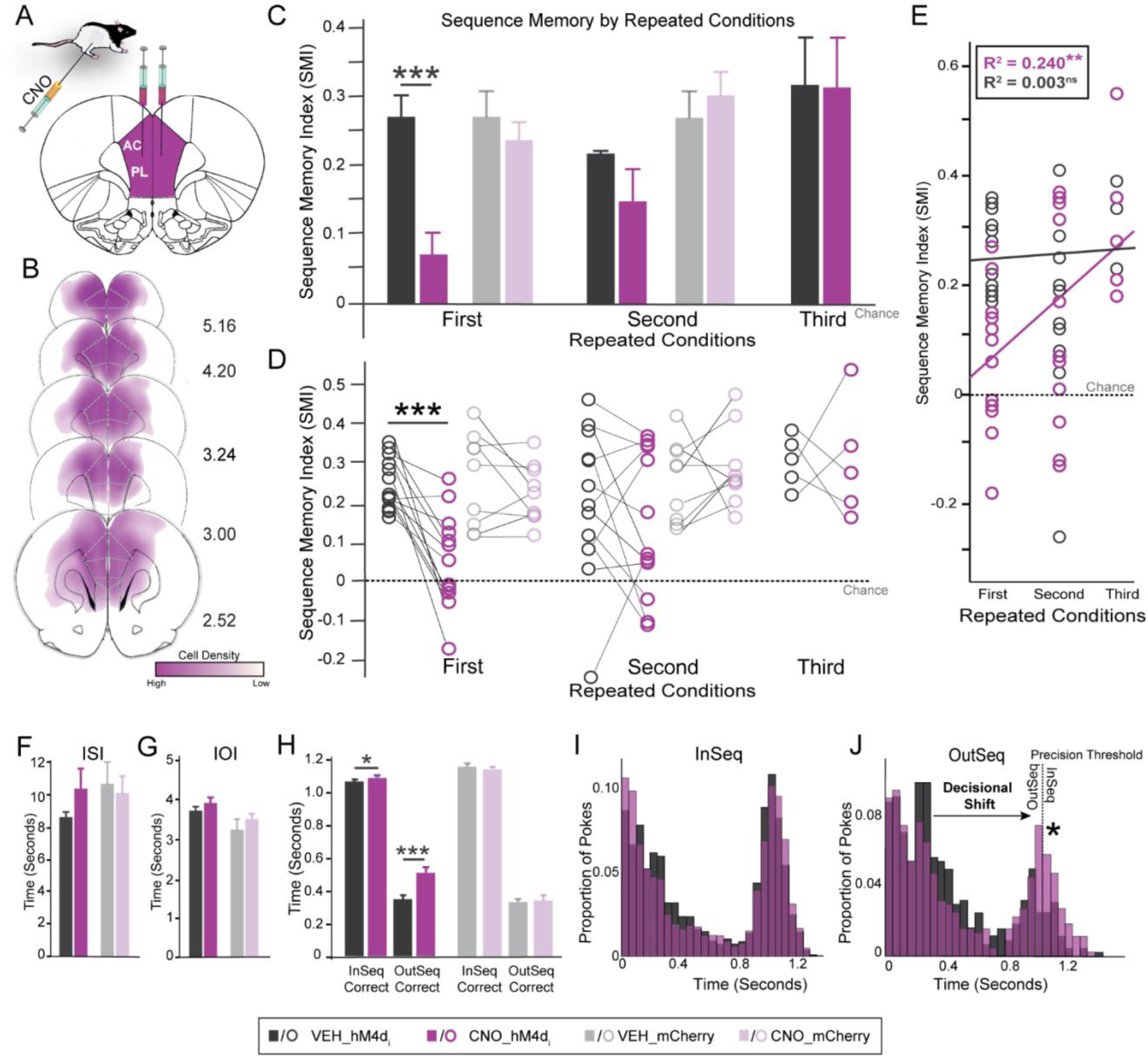
PL cortex is needed for sequence memory. Suppressing PL activity in the hM4Di+ group impaired sequence memory, but there were no significant effects with vehicle administration or in the mCherry-only group (CNO and vehicle). (A) AAV9.hM4Di was injected into PL (bilaterally). Systemic CNO administration (i.p. injection) was used to suppress PL activity. (B) Schematic representation of AAV9.hM4Di viral spread in PL for all rats (n =13). Numbers to the right of each section indicates distance (mm) anterior to bregma according to Paxinos and Watson (2004). (C) The first repeated condition for the hM4Di+ group was significantly different between vehicle and CNO injection. The second and third repeated condition did not show any significant differences. Moreover, the mCherry-only group did not show any significant differences between vehicle and CNO injections across all repeated conditions. Since the third repeated condition in the hM4Di+ group did not show any significant differences between vehicle and CNO, a third repeated condition was not run for the mCherry-only group. (D) Each individual rat’s performance per repeated condition for both hM4Di+ and mCherry-only groups. (E) In the hM4Di+ group, there was a moderate positive linear relationship between repeated conditions and CNO injections and no significant relationship between repeated conditions and vehicle injections. (F-H) We tested if motor behaviors such as running and poking were affected under all conditions. We first looked at inter-sequence-interval (F) and inter-odor-interval (G) for overall activity levels in the task. (F) inter-sequence-interval is the amount of time (s) it took the rat to run between sequences. There was no significant difference between vehicle and CNO injection for both hM4Di+ and mCherry-only groups. (F) Inter-odor-interval is the amount of time (s) spent between odor trials. There was no significant difference between vehicle and CNO injection for both the hM4Di+ and mCherry-only groups. (G) We next examined poke times for InSeq_correct_ and OutSeq_correct_ trials. In the hM4Di+ group, there was a significant difference between vehicle and CNO for both InSeq_correct_ and OutSeq_correct_ poke times. However, the mCherry-only group did not show any significant differences for InSeq_correct_ and OutSeq_correct_. (I) A histogram of the hM4Di+ group poke times on All InSeq trials shows only subtle shifts in behavior where CNO (magenta) caused a slight decrease in the proportion of pokes near 0.3s-0.4s. (J) A histogram of the hM4Di+ group poke times on All OutSeq trials shows a clear shift (from left to right, indicated with a star) between vehicle (black) and CNO (magenta), signifying a decisional shift where the rat incorrectly identified OutSeq odors as InSeq. Abbreviations: CNO, clozapine-n-oxide; VEH, vehicle; InSeq, in sequence item(s); OutSeq, out of sequence item(s); ISI, inter-sequence-interval; IOI, inter-odor-interval; SMI, sequence memory index; PL, prelimbic cortex; AC, anterior cingulate cortex; * = p < 0.05; ** p < 0.01; *** = p < 0.001.

We also examined the immunolabeling of mCherry terminals, and found the pattern of fiber expression was characteristic of medial prefrontal cortex projection sites to the forebrain including the orbital and insular cortices, ventral striatum, PER and entorhinal cortices (Figure 5C), and the entire intralaminar and midline thalamus including predominantly RE (Figure 4C), indicative of good anterograde transport, allowing for CNO inhibition of PL terminal fibers to RE (Vertes, 2002; Hoover and Vertes, 2007; Vertes et al., 2007; Hoover and Vertes, 2012) and PER (Furtak et al., 2007). The placement of cannula for intracerebral infusions was confirmed for all rats using cresyl violet stain, and their locations are mapped in Figure 4B, 5B, and S5.

**Figure 4.**
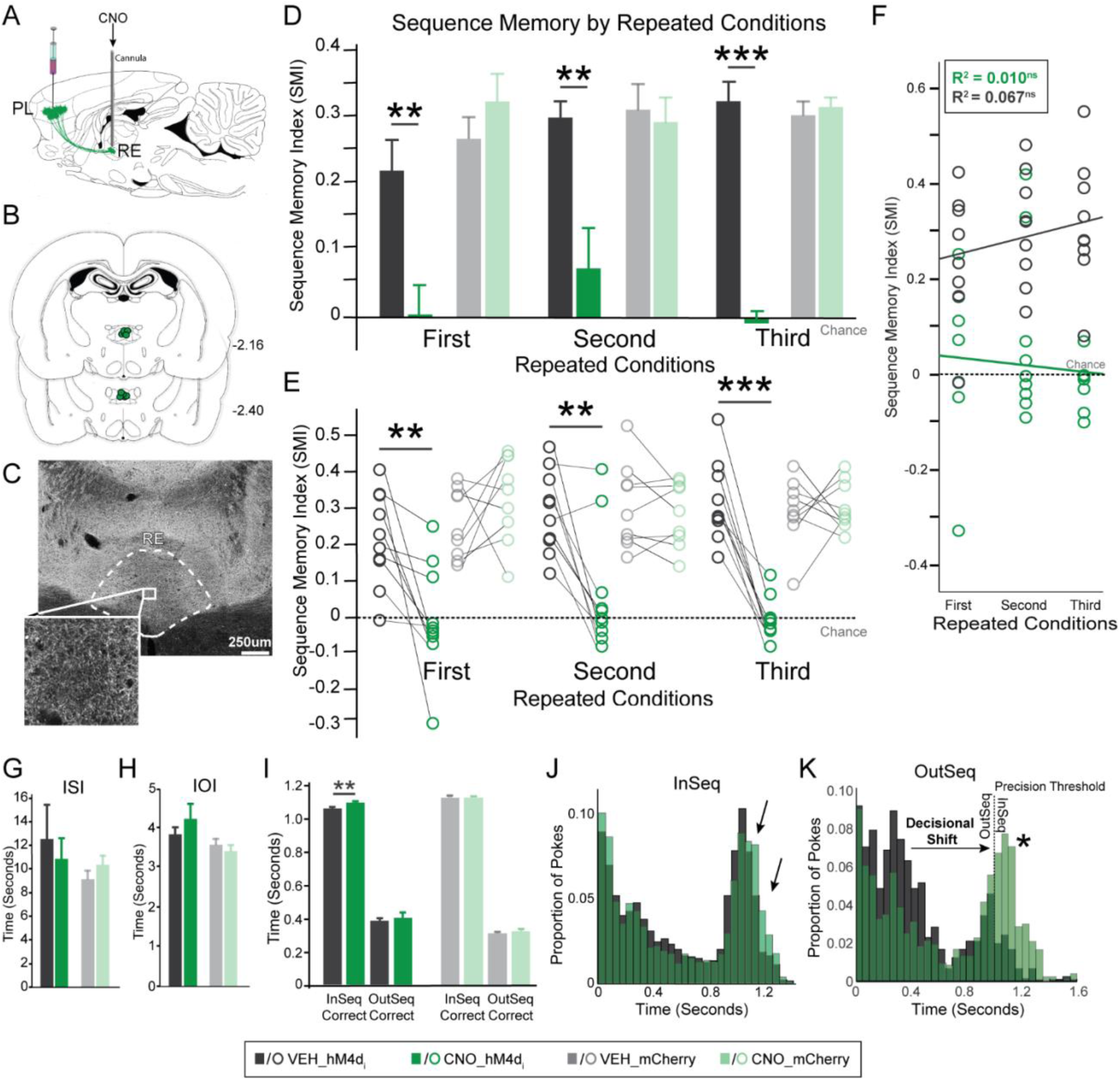
Synaptic silencing of the PL➔ RE pathway eliminated sequence memory. PL➔RE silencing in the hM4Di+ group eliminated sequence memory while vehicle and the mCherry-only group did not show any significant effects on sequence memory. (A) Guide cannula were implanted targeting RE, such that CNO infusions would inactivate PL terminals in RE. (B) Microinfusion injector tips targeting RE were located and represented (green circles) for all rats (n=10). Numbers to the right of each section indicates distance (mm) posterior to bregma according to Paxinos and Watson (2004). (C) Coronal image of the thalamus from a representative rat shows the AAV9.hM4Di viral construct expressed in RE axons. (D) All three repeated conditions for the hM4Di+ group showed significant differences between vehicle and CNO infusions. The mCherry-only group did not show any significant differences between vehicle and CNO infusions. (E) Shows individual rat’s performance per repeated conditions for both hM4Di+ and mCherry-only groups. (F) In the hM4Di+ group, there was not significant relationship between repeated conditions and infusions (vehicle and CNO). (G-I) We examined if any motor behaviors such as running and poking were affected with CNO infusions. (G) In terms of inter-sequence-interval there was no significant difference between RE vehicle and CNO infusions for both hM4Di+ and mCherry-only groups. (H) With regards to inter-odor-interval, there was no significant difference between RE vehicle and CNO infusions for both hM4Di+ and mCherry-only groups. (I) In the hM4Di+ group, there was a significant difference between vehicle and CNO for InSeq_correct_ poke times. There was however, no significant differences between vehicle and CNO for OutSeq_correct_ pokes times. Moreover, there was no significant difference between vehicle and CNO for InSeq_correct_ and OutSeq_correct_ poke times for the mCherry-only group. (J) A histogram of the hM4Di+ group poke times for All InSeq trials remained relatively similar. CNO (green) caused a slight decrease in the proportion of pokes near 0.35s-0.6s, and an increase near the 1s decision threshold indicated by the arrows. (K) A histogram of the hM4Di+ group poke times on All OutSeq trials shows a clear shift (from left to right; indicated with a star) between vehicle (black) and CNO (green), signifying a decisional shift where the rat incorrectly identified OutSeq odors as InSeq. Abbreviations: CNO, clozapine-n-oxide; VEH, vehicle; InSeq, in sequence item(s); OutSeq, out of sequence item(s); ISI, inter-sequence-interval; IOI, inter-odor-interval; RE, nucleus reuniens of the thalamus; PL, prelimbic cortex; SMI, sequence memory index; * = p < 0.05; ** p < 0.01; *** = p < 0.001.

**Figure 5.**
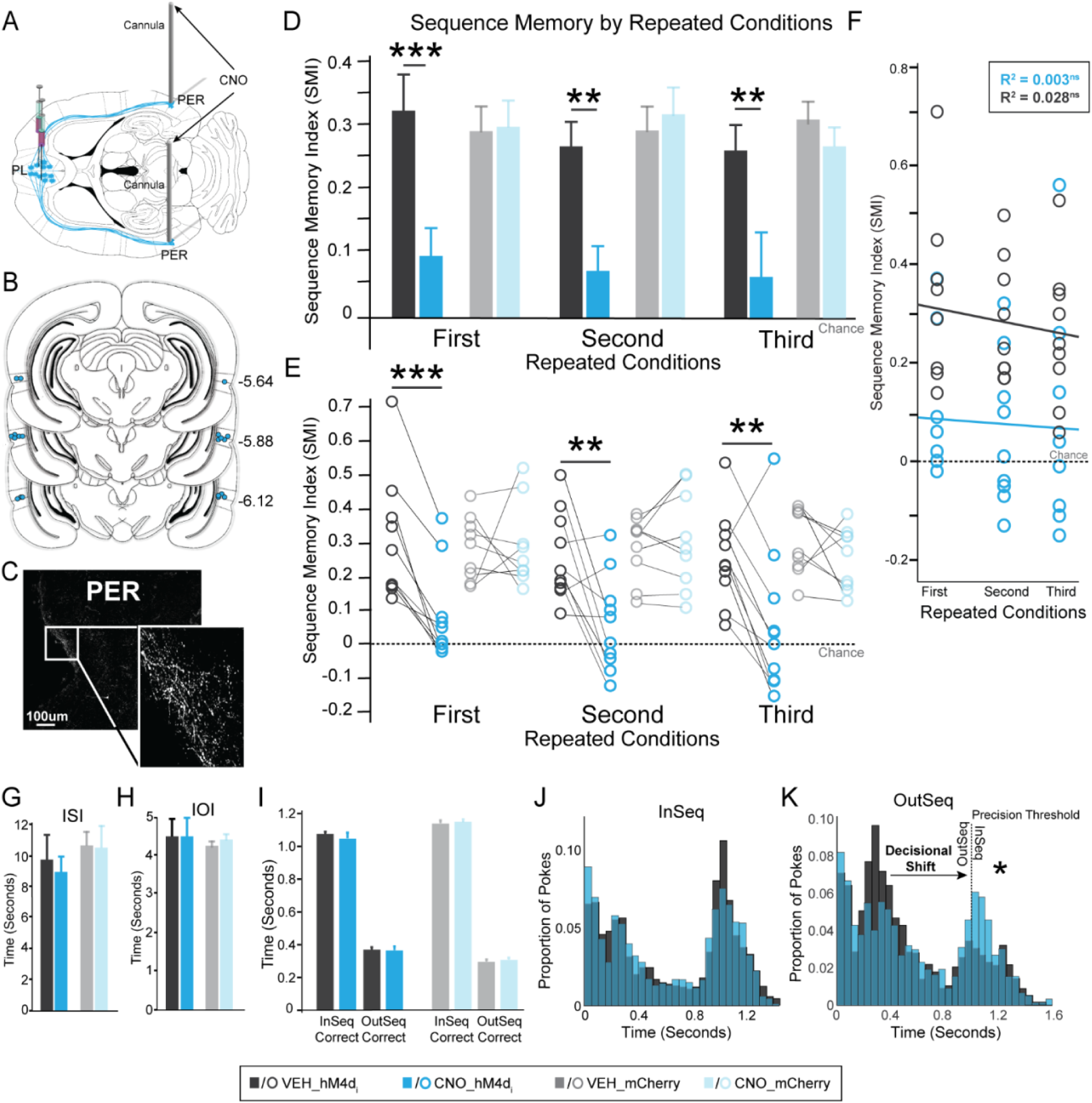
Synaptic silencing of the PL➔PER pathway eliminated sequence memory. PL ➔ PER silencing in the hM4Di+ group eliminated sequence memory while vehicle and the mCherry-only group did not show any significant effects on sequence memory. (A) Guide cannula were implanted targeting PER, such that CNO infusions would inactivate PL terminals in PER. (B) Microinfusion injector tips targeting PER were located and represented (blue circles) for all rats (n=10). Numbers to the right of each section indicates distance (mm) posterior to bregma according to Paxinos and Watson (2004 (C)). (D) Coronal image of the thalamus from a representative rat shows the AAV9.hM4Di viral construct expressed in PER axons. (E) All three repeated conditions for the hM4Di+ group showed significant differences between vehicle and CNO infusions. The mCherry-only group did not show any significant differences between vehicle and CNO infusions. (F) Shows individual rat’s performance per repeated conditions for both hM4Di+ and mCherry-only groups. (F) In the hM4Di+ group, there was not significant relationship between repeated conditions and infusions (vehicle and CNO). (G-I) We examined if any motor behaviors such as running and poking were affected with CNO infusions. (G) In terms of inter-sequence-interval there was no significant difference between PER vehicle and CNO infusions for both hM4Di+ and mCherry-only groups. (H) With regards to inter-odor-interval, there was no significant difference between PER vehicle and CNO infusions for both hM4Di+ and mCherry-only groups. (I) In the hM4Di+ and mCherry-only groups there was no significant difference between vehicle and CNO for InSeq_correct_ and OutSeq_correct_ poke times. (J) A histogram of the hM4Di+ group poke times on All InSeq trials show no obvious shifts in poking behavior. (k) A histogram of the hM4Di+ group poke times on All OutSeq trials shows a clear shift (from left to right; indicated with a star) between vehicle (black) and CNO (blue), signifying a decisional shift where the rat incorrectly identified OutSeq odors as InSeq. Abbreviations: CNO, clozapine-n-oxide; VEH, vehicle; InSeq, in sequence item(s); OutSeq, out of sequence item(s); ISI, inter-sequence-interval; IOI, inter-odor-interval; PER, perirhinal cortex; PL, prelimbic cortex; SMI, sequence memory index; * = p < 0.05; ** p < 0.01; *** = p < 0.001.

We also mapped the spread of mCherry cells for the mCherry-only control rats (Figure S5A). While infected cells in mCherry-only rats were visualized outside of the medial wall of the prefrontal cortex, extending to the orbital and motor cortices, the relative density and pattern of labeling of terminal fibers to target sites (RE and PER) was similar to hM4Di+ rats Furthermore, the null behavioral effects of these rats allowed us to confirm our findings were not associated with non-specific effects related to the viral construct or CNO. Characteristic mCherry expression in PL and anterior cingulate cortex from a representative case of both hM4Di+ and mCherry-only rat is depicted in Figure S5. Lastly, while labeling of neurons was virtually exclusive to the injection sites, sporadic cells were seen in target sites, (less than ∼ 0.1-1%), most likely associated with weak retrograde transport of adenovirus constructs (Castle et al., 2014; Tervo et al., 2016) observed in other hM4Di applications (DiBenedictis et al., 2015).

### PL cortex is critical to sequence memory

Evidence suggests that PL makes essential contributions to sequence memory (DeVito and Eichenbaum, 2011). Here, we examined the contribution of PL in sequence memory via systemic CNO (i.p., 1mg/kg) or vehicle injections. We refer to each corresponding vehicle and CNO injection as a repeated condition across behavioral sessions. We observed that the suppression of PL neurons with CNO in the hM4Di+ group (SMI: 0.056 ± 0.121) significantly reduced SMI scores compared to vehicle injection (SMI: 0.254 ± 0.062) the first time we ran this condition (Figure 3C and 3D; first: t_(12)_ = 5.200, p = 2.210×10^-4^, Cohen’s d = 1.449), however, the second and third repeats of this condition was not effective (second: t_(12)_ = 1.039, p = 0.325; third: t_(4)_ = 0.031, p = 0.970). Overall, we performed three repeated conditions for the first cohort of hM4Di+ (n=5). We reduced to two repeated conditions for the second cohort of hM4Di+ (n=8) and mCherry-only (n=9), since the third repeated condition did not have an effect after CNO administration. In order to see if there was a relationship between CNO administration and repeated conditions, we performed a Pearson’s correlation. Figure 3E shows that there was a moderate positive linear relationship between the repeated conditions and CNO in the hM4Di+ group (Pearson’s r = 0.490, R^2^ = 0.240, p = 0.003), but no relationship between the repeated condition and vehicle in the hM4Di+ group (Pearson’s r = 0.057, R^2^ = 0.003, p = 0.380). An ANOVA yielded similar results showing a significant relationship between repeated conditions and SMI for CNO injections (F_(1,29)_ = 9.167, p = 0.005). Importantly, CNO administration had no significant effects in the mCherry-only group (Figure 3C and 3D; first: t_(8)_ = 1.208, p = 0.262; second: t_(8)_ = −0.623, p = 0.551), indicating the observed effects in the hM4Di+ group are specific to hM4Di receptor activity in PL neurons. Lastly, sequence memory performance levels were similar between sequence1 and sequence2, with no significant differences between sequences (Figure S3B; first: vehicle, t_(11)_ = −1.023, p = 0.328, CNO, t_(11)_ = −1.231, p = 0.244; second: vehicle, t_(9)_ = 2.086, p = 0.067, CNO, t_(10)_ = 0.533, p = 0.606), suggesting a general sequence memory deficit.

Next, we examined the possibility that hM4Di suppression of PL activity produced non-mnemonic effects relevant to the sequence task. To test for this possibility, we measured the time it took to run between sequences (inter-sequence-interval), the time spent between each odor trial (inter-odor-interval), and nose poke times. There were no significant differences between vehicle and CNO injections in either the hM4Di+ or mCherry-only groups in the inter-sequence-interval (Figure 3F; hM4Di+: t_(25)_ = −1.427, p = 0.166; mCherry-only: t_(17)_ = 0.454, p = 0.655), suggesting rats ran between sequences at similar rates. Nor did we find effects on the inter-odor-interval (Figure 3G; hM4Di+: t_(25)_ = −1.616, p = 0.119; mCherry-only: t_(17)_ = −1.056, p = 0.306), suggesting rats collected water rewards and engaged odors at similar rates. We looked at whether they had any holding bias during the task between vehicle and CNO and found there was no significant difference between holding (>1s; t_(12)_ = −0.875, p = 0.399) and not holding (<1s; t_(12)_ = 0.506, p = 0.622). We also looked at whether CNO had an effect on nose poke times on InSeq_correct_ and OutSeq_correct_ trials (Figure 3H). In the hM4Di+ group, CNO suppression of PL activity significantly increased poke times in both InSeq_correct_ and OutSeq_correct_ trials (hM4Di+: InSeq_Correct_, t_(25)_ = - 2.672, p = 0.013, OutSeq_Correct_, t_(25)_ = −3.944, p = 0.001). However, there were no significant differences in the mCherry-only group (mCherry-only: InSeq_Correct_, t_(17)_ = 0.973, p = 0.342, OutSeq_Correct_, t_(17)_ = −0.186, p = 0.853) suggesting i.p. CNO itself does not affect nose poke behavior in the task. The increase in hold times in the hM4Di+ group might indicate uncertainty (knowing whether a trial was InSeq or OutSeq) and decisional differences, rather than deficits related to basic poke and hold behavior. In order to visualize and further examine this possibility, we looked at detailed poke distributions for both InSeq and OutSeq trials (Figure 3I and 3J). The InSeq distribution shows that the proportion of pokes remained relatively similar between vehicle and CNO in the hM4Di+ group, with only modest differences. However, on OutSeq trials, there is a clear shift in the proportion of trials near the 1s decision threshold indicative of inaccurately making InSeq decisions. Thus, the nose poke differences indicate a decisional shift in the hM4Di+ group following CNO administration. Overall, these results demonstrate that suppression of PL neurons (via systemic CNO administration) in the hM4Di+ group impaired memory for sequences of events but the effect decreases with subsequent administrations of CNO.

### Synaptic silencing PL➔RE projections eliminated sequence memory

Our primary goal was to examine whether specific PL inputs to RE and PER, structures heavily interconnected with the hippocampus, contribute to sequence memory (Eichenbaum, 2017b). We tested top down PL inputs by using intracranial CNO infusions (1μl at a 1μg/μl concentration per cannula) targeting RE and PER (within subject) on different days. The daily schedule for RE and PER infusions were randomized and counterbalanced (Figure 1F) across rats and repeated conditions to avoid order effect.

We first examined PL➔RE projections (Figure 4A). We found that silencing of PL terminals in RE in the hM4Di+ group significantly impaired sequence memory (Figure 4D and 4E). There was a clear difference between CNO and vehicle infusions (F_(1,9)_ = 130.850, p = 1.000×10^-6^), however, there were no differences across repeated conditions (Figure 4D and 4E; F_(1.479,13.312)_ = 1.012, p = 0.366). Thus, silencing PL➔RE synapses powerfully and repeatedly eliminated sequence memory. The mCherry-only group showed no significant differences between CNO and vehicle infusions and no differences across repeated condition (Figure 4D and 4E; infusion: F_(1,8)_ = 0.492, p = 0.503; repeated conditions: F_(1.818,14.545)_ = 0.025, p = 0.967), controlling for non-specific CNO effects in RE. A simple linear regression was carried out to investigate the relationship between SMI (CNO and vehicle) and repeated condition for the hM4Di+group. There was no significant relationship between the repeated condition and SMI following either CNO or vehicle infusions (Figure 4F; CNO: Pearson’s r = 0.101, R^2^ = 0.010, p = 0.307; vehicle: Pearson’s r = 0.259, R^2^ = 0.067, p = 0.096). In the hM4Di+ group, sequence1 and sequence2 were similar across all repeated conditions (Figure S3C; first: vehicle, t_(9)_ = 0.989, p = 0.348, CNO, t_(9)_ = −1.665, p =.130; second: vehicle, t_(9)_ = 0.242, p = 0.814, CNO, t_(9)_ = 0.754, p = 0.322, third: vehicle, t_(9)_ = −0.506, p = 0.625, CNO, t_(9)_= 1.069, p = 0.316), indicating a general sequence memory deficit.

We looked at the non-mnemonic effects of PL➔RE silencing examining inter-sequence-interval, inter-odor-interval, and nose poke behavior. There were no significant differences between vehicle and CNO injections in either the hM4Di+ or mCherry-only groups in the inter-sequence-interval, nor with the inter-odor-interval (Figure 4G and 4H; hM4Di+: inter-sequence-interval, t_(29)_ = 0.502, p = 0.611, inter-odor-interval, t_(29)_ = 0.394, p = 0.697; mCherry: inter-sequence-interval, t_(26)_ = −1.123, p = 0.272, inter-odor-interval, t_(26)_ = 1.119, p = 0.273). There was no significant holding bias during the task between vehicle and CNO (holding, >1s: t_(29)_ = −0.639, p = 0.528; not holding, <1s: t_(29)_ = −0.056, p = 0.956). Additionally, nose poke times were mostly similar between vehicle and CNO conditions with both the hM4Di+ and mCherry-only groups following RE infusions (Figure 4I; hM4Di+: OutSeq_Correct_, t_(29)_ = −0.709, p = 0.485, mCherry-only: InSeq_Correct_, t_(26)_ = −0.020, p = 0.982, OutSeq_Correct_, t_(26)_ = −0.777, p = 0.448), but there was a slight and significant increase in the amount of time the rat held during the CNO condition in the hM4Di+ group on InSeq trial (InSeq_Correct_: t_(29)_ = −2.760, p = 0.011). We followed up with this analysis and looked at the detailed poke distributions in the hM4Di+ group for all InSeq and OutSeq trials (Figure 4J and 4K). The InSeq distribution shows that the proportion of pokes remained relatively similar between vehicle and CNO in the hM4Di+ group, with a slight increase in holding shown in Figure 4J. As shown in Figure 4K, CNO in the hM4Di+ group caused a decrease in the proportion of pokes near the short distribution peak (∼0.35-0.6s), and an increase near the 1s decision threshold, compared to vehicle infusions. The nose poke differences indicate a decisional shift in the hM4Di+ group following CNO infusions. Overall, these results provide strong evidence that silencing PL➔RE leads to inaccurate decisions, not a basic deficit in poke and hold behavior, reflecting deficits in sequence memory.

### Synaptic silencing PL➔PER projections eliminated sequence memory

We next looked at PL➔PER projections (Figure 5A). Silencing the PL➔PER projections significantly and consistently impaired sequence memory across repeated conditions in the hM4Di+ group (Figure 5D and 5E; infusion: F_(1,8)_ = 62.750, p = 4.700×10^-5^; repeated condition: F_(1.592,12.733)_ = 1.466, p = 0.260). A linear regression showed no significant relationship between repeated conditions and SMI following PER infusions with either CNO or vehicle in the hM4Di+ group (Figure 5F; CNO: Pearson’s r = 0.053, R^2^ = 0.003, p = 0.399; vehicle: Pearson’s r = 0.167, R^2^ = 0.028, p = 0.202). Furthermore, CNO did not have a significant effect on sequence memory in the mCherry-only group compared to vehicle infusions (Figure 5D and 5E; infusion: F_(1,8)_ = 0.690, p = 0.430; repeated condition: F_(1.715,13.722)_ = 0.092, p = 0.886). Additionally, there was no significant difference between the two sequences under any conditions (Figure S3D; first: vehicle, t_(9)_ = 0.013, p = 0.990, CNO, t_(8)_ = −0.928, p =.380; second: vehicle, t_(9)_ = −0.169, p = 0.870, CNO, t_(9)_ = −1.101, p = 0.300; third: vehicle, t_(9)_ = 1.003, p = 0.342, CNO, t_(9)_ = 0.630, p = 0.545), thus indicating a general sequence memory deficit.

We looked at the non-mnemonic effects of PL➔PER silencing examining inter-sequence-interval, inter-odor-interval, and nose poke behavior. There were no significant differences between vehicle and CNO injections in either the hM4Di+ or mCherry-only groups in the inter-sequence-interval, nor with the inter-odor-interval following PER infusions (Figure 5G and 5H; hM4Di+: inter-sequence-interval, t_(29)_ = 0.634, p = 0.534, inter-odor-interval, t_(29)_ = 0.456, p = 0.658; mCherry: inter-sequence-interval, t_(26)_ = 0.082, p = 0.935, inter-odor-interval, t_(26)_ = −0.874, p = 0.390). Furthermore, there was no significant holding bias during the task between vehicle and CNO (holding: t_(29)_ = 0.322, p = 0.750; not holding: t_(29)_ = 1.595, p = 0.122). Additionally, nose poke times were similar between vehicle and CNO conditions with hM4Di+ and mCherry-only groups following PER infusions (Figure 5I; hM4Di: InSeq_Correct_: t_(29)_ = 0.621, p = 0.540, hM4Di_+_: OutSeq_Correct_: t_(29)_ = 0.055, p = 0.956, mCherry-only: InSeq_Correct_: t_(26)_ = −0.457, p = 0.652, mCherry-only: OutSeq_Correct_: t_(26)_ = −0.707, p = 0.486). We followed up with this analysis and looked at the detailed poke distributions in the hM4Di+ group for all InSeq and OutSeq trials following PER infusions (Figure 5J and 5K). The proportion of pokes in the InSeq distribution were similar between vehicle and CNO in the hM4Di+ group (Figure 5J). As shown in Figure 5K, CNO caused a decrease in the proportion of pokes short poke distribution, (∼.20-.4s) and an increase near the 1s decision threshold, compared to vehicle infusions. Thus, the nose poke differences indicate a decisional shift in the hM4Di+ group following CNO administration. Similar to PL➔RE, these results provide strong evidence that silencing PL➔PER leads to inaccurate decisions, not a basic deficit in poke and hold behavior, reflecting deficits in sequence memory.

### Differential Role of PL Top down inputs to RE and PER

The results demonstrate that activity in both PL➔ RE and PL➔PER projections are essential to sequence memory. Next, we directly compared the effects of silencing PL➔RE and PL➔PER projections. Overall, silencing of PL➔RE and PL➔PER projections were not significantly different from each other across the repeated conditions in the hM4Di+ group (region: F_(1,8)_ =1.487, p = 0.257; repeated condition: F_(1.340,10.720)_ = 0.291, p = 0.667; repeated condition X region: F_(1.331, 10.647)_ = 0.343, p = 0.632). We then examined whether silencing PL➔RE and PL➔PER projections impaired different memory retrieval strategies that support sequence memory. Our approach was based on the conceptual model shown in Figure 6A (see Reeders et al., 2018). This model illustrates theoretical performance curves that would be obtained when using working memory and temporal context memory retrieval strategies plotted as a function of sequential lag distances on OutSeq items. With a working memory strategy repeated items would be easier to detect at shorter lags because these items occur more recently. Conversely, with a temporal context memory strategy, repeated items would be easier to detect at longer lags because these items are further away in the original sequence. Therefore, we examined the performance of the OutSeq probe trials across lags, focusing on repeated items (also called backward lags) in order to test the contributions of temporal context memory and working memory. For this analysis, we calculated the percent change in performance (CNO-vehicle) for each rat on each lag.

**Figure 6.**
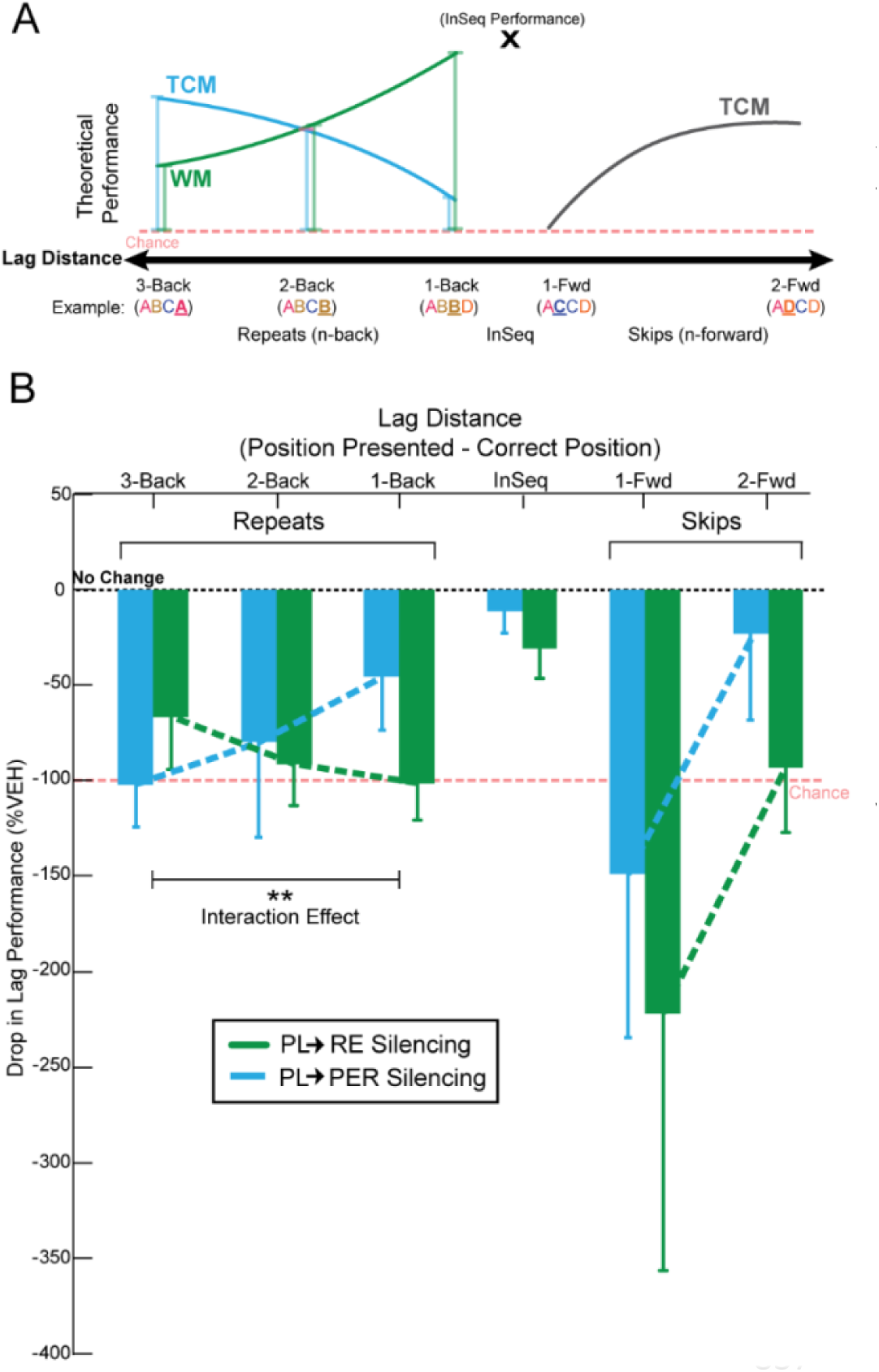
PL➔RE pathway supports a working memory retrieval strategy, while PL➔PER pathway supports a temporal context memory retrieval strategy. We used a lag analysis in order to evaluate performance across items that skipped ahead in the sequence (lags = +1, +2) and items that repeated (lags = −3, −2, −1). (A) A conceptual model based on temporal context memory and working memory across different lags (see also Reeders et al., 2018). Temporal context memory (blue and gray) predicts the retrieval probability of items decreases as the lag distance increases. Working memory (green) predicts recency for strength of memory for repeated items only. InSeq performance is represented with an X. (B) We tested contributions of PL➔RE and PL➔PER pathways by analyzing the percent change in performance (CNO minus vehicle). As the backward lag distance increased on repeated items, PL➔RE silencing resulted in decreased performance consistent with a loss of working memory, while PL➔PER silencing resulted in increased performance consistent with a loss in temporal context memory. This pattern of impairment showed a strong pathway x behavior interaction effect. However, we did not find any significant main effects between regions or lags in the backward or forward direction. Abbreviations: InSeq, in sequence item(s); RE, nucleus reuniens of the thalamus; PER, perirhinal cortex; PL, prelimbic cortex; TCM, temporal context memory; WM, working memory; ** = p < 0.01.

We first looked at items that were repeated in a sequence to see if there was any difference in the impairment patterns dependent on silencing PL➔RE and PL➔PER projections. A one-sample t-test was run to test for impairments on repeated items. PL➔RE silencing resulted in significant differences from no change in performance (0%) on all backward lags (3-back, t_(14)_ = −2.388, p = 0.016; 2-back, t_(26)_ = −4.252, p = 1.21 x 10^-4^; 1-back, t_(29)_ = −5.274, p = 6.00 x 10^-6^). Additionally, PL➔PER silencing resulted in a significant difference on the 3-back, and showed trends toward significant differences from no change in performance on the 2-back and 1-back lags (3-back, t_(21)_ = −4.579, p = 8.15 x 10^-5^; 2-back, t_(27)_ = −1.609, p = 0.059; 1-back, t_(28)_ = −1.639, p = 0.056). These results indicate that repeated items were affected by silencing both PL➔RE and PL➔PER pathways. However, the important question is whether there was a difference in the performance patterns across lags when directly comparing the effects of silencing PL➔RE and PL➔PER pathways. A repeated-measures ANOVA was run revealing a significant interaction effect between PL➔RE silencing and PL➔PER silencing (region X backward: F_(1.968, 23.619)_ = 5.395, p = 0.012), with a large effect size (η_p_^2^ = 0.310; Cohen, 1973). In the hM4DI+ group, CNO infusions into RE had the largest effect on 1-back, then 2-back, and the smallest effect on 3-back (Figure 6B). In the hM4DI+ group, CNO infusions into PER had the opposite pattern, with the largest effect on 3-back, then 2-back, and the smallest effect on 1-back (Figure 6B). These impairment patterns match the expected performance decrements (drop lines in Figure 6A) from a selective loss of working memory following PL➔RE silencing, and a selective loss in temporal context memory following PL➔PER silencing.

Next, performance for InSeq trials (lag = 0) was tested against no change in performance (0%) following PL➔RE and PL➔PER silencing. There were no significant differences (InSeq_RE_: t_(29)_ = −1.189, p = 0.122; InSeq_PER_: t_(28)_ = −1.004, p = 0.162). The suggests that the InSeq trials were not clearly affected by silencing PL➔RE and PL➔PER pathways. We found no significant differences comparing PL➔RE and PL➔PER on InSeq performance levels using a paired-samples t-test (t_(28)_ = −0.982, p = 0.334).

Lastly, we looked at if performance on items that skipped ahead in the sequence (forward lags) were affected by PL➔RE and PL➔PER silencing. We found that PL➔RE silencing had a trend towards significance on 1-forward (t_(29)_ = −1.644, p =.055), and a significant difference on 2-forward (t_(29)_ = −2.725, p = 0.006). PL➔PER silencing showed a significant difference on the 1-forward (t_(28)_ = 1.732, p = 0.047) and no significant difference on the 2-forward (t_(27)_ = −0.524, p = 0.303). Overall, this suggests that performance levels on items that skipped ahead in the sequence were affected by silencing PL➔RE and PL➔PER pathways. Again, the important question is whether there was a difference in the patterns when directly comparing the silencing effects in the PL➔RE and PL➔PER pathways. We used a repeated-measures ANOVA and did not find any significant interaction effects in the forward direction (region X forward: F_(1,27)_ = −.012, p = 0.915).

## Discussion

### Summary of main findings

The present study examined the hypothesis that top down prefrontal projections contribute to sequence memory, and that separate projections control the selection of different retrieval strategies. First, we showed that PL is critical to sequence memory by suppressing PL activity, thereby impairing sequence memory; this is consistent with other reports (e.g., Hannesson et al., 2004; DeVito and Eichenbaum, 2011). However, this result alone does not speak to a role for PL in top down control of sequence memory. Therefore, we directly manipulated PL circuitry using an hM4Di synaptic silencing approach. We found that suppressing activity in the PL➔RE or PL➔PER pathway effectively eliminated sequence memory. These results unambiguously demonstrate that top down PL projections are essential to sequence memory. Lastly, we used a detailed behavioral lag analysis to determine the differential roles of the PL➔RE or PL➔PER pathway. We found that silencing the PL➔RE pathway disrupted backward lags with a pattern resembling a loss of working memory, whereas silencing the PL➔PER pathway disrupted backward lags with a pattern resembling a loss of temporal context memory. Theoretically, working memory and temporal context memory (i.e., graded retrieval strength based on temporal proximity) differentially contribute when items are repeated (see also Reeders et al., 2018). That is, working memory strategies lead to better performance on shorter lags (e.g., ABC<u>C</u>, lag = −1) which was disrupted when suppressing PL➔RE activity, and temporal context memory strategies lead to better performance at longer lags (e.g., ABC<u>A</u>, lag = −3) which was disrupted when suppressing PL➔PER activity in the backward direction. This strong pathway-specific interaction effect demonstrates, for the first time, that top down PL projections control sequence memory, and suggests that RE and PER pathways provide mechanisms enabling the regulation of ongoing retrieval strategies.

### Testing PL projections with DREADDs in sequence memory

In order to test the top down role of PL in sequence memory we suppressed synaptic activity in specific PL projections (Sesack et al., 1989; Chiba et al., 2001; Vertes, 2002; Hoover and Vertes, 2007) using Gi-coupled (hM4Di) DREADDs (Mahler et al., 2014; Roth, 2016) combined with multiple chronic cannula in rats. Briefly, activation of hM4Di in presynaptic terminals reduces transmission and effectively silences the affected projection pathways (Stachniak et al., 2014; Lichtenberg et al., 2017). Here the focus was on terminal fields in RE (Sesack et al., 1989; Chiba et al., 2001; Vertes, 2002; Hoover and Vertes, 2007; Vertes et al., 2015) and PER (Furtak et al., 2007) because both serve as critical links supporting bidirectional communication between the hippocampus and medial prefrontal cortex (e.g., Allen et al., 2013; Vertes et al., 2015; Eichenbaum, 2017b). To ensure our manipulations localized to these pathways we carefully mapped the PL hM4Di expression areas, visualized PL➔RE and PL➔PER projection terminals via the co-expression of mCherry (enhanced with IHC), and mapped the tip locations of the infusion cannula within RE and PER.

An important consideration in using hM4Di for brain-behavior relationships is controlling for non-specific CNO or infusion effects (Smith et al., 2016; Gomez et al., 2017). Thus, we used a fully-crossed 2 (hM4Di and hM4Di-free) x 2 (CNO and vehicle) experimental design. Notably, the only effects observed in any of our manipulations arose when we activated hM4Di with CNO and tested sequence memory. We also ran several analyses testing the alternative hypothesis that non-memory related behavioral effects could account for the reductions in performance in the sequence memory task. We did not observe any effects (under any conditions) on the time it took rats to run between sequences, on odor sampling and reward retrieval activity, nor on the overall frequency of nose pokes in which rats held for >1s or less <1s (analyzed by ignoring the sequential status of items; see Results). In fact, a detailed analysis of all nose poke times under hM4Di+ and CNO conditions showed a pattern resembling a shift toward inaccurate OutSeq decisions. This was most obvious when plotting poke time histograms and observing an increase in the overall frequency of holds on OutSeq items, without large changes in distribution variability or peak times. In fact, we would have expected increased premature and variable responses following PL inactivation if the rats had shifted to simple reaction time behaviors (Narayanan et al., 2006). Lastly, the sequence memory was consistent across two different sequences. This is important because it eliminates the possibility that rats held a single sequence in working memory focusing on a single strategy throughout the entirety of a session, and instead forced rats to repeatedly retrieve sequences from longer term memory stores.

### PL pathways to RE and PER control retrieval strategies in sequence memory

Generally, it is thought that a major role of the medial prefrontal cortex (including PL) is to control memory retrieval strategies (Shimamura et al., 1995; Dobbins et al., 2002; Euston et al., 2012; Preston and Eichenbaum, 2013; Jadhav et al., 2016; Eichenbaum, 2017b). We tested this in specific projection pathways during memory for sequences of events. Importantly, the sequence task we used has been related to episodic-like memory processing and depends on the use of multiple cognitive strategies for optimal performance (Allen et al., 2014; 2015). Using this task, we recently presented evidence using with BOLD fMRI in humans supporting a theoretical model in which working memory and temporal context memory (Howard and Kahana, 2002; Roberts et al., 2014; Kragel et al., 2015) differentially support performance on items that are repeated at across lags (e.g., ABC<u>C</u>, lag = −1; ABC<u>A</u>, lag = −3) (see Reeders et al., 2018). If rats were using a working memory strategy, then repeated items would be easiest to detect at short lags because those items occurred more recently. However, if rats were using a temporal context memory strategy (i.e., graded retrieval probabilities determined by the temporal proximity of items in the sequence), then repeated items would be the easiest to detect with longer lags, because they occurred with the least temporal proximity. Here, incidental retrieval of nearby items interferes with OutSeq determinations. Thus, the sequence task places pressure on the ability of rats to regulate retrieval strategies at different lag distances for optimal performance using working memory on shorter lags, and temporal context memory on longer lags. We found that suppressing PL➔RE activity had the largest effect on lags = −1 and the smallest effect on lags = −3. This pattern resembles a reduction in a working memory retrieval strategy and is consistent with the role of RE in spatial working memory tasks (Cassel et al., 2013; Griffin, 2015; Vertes et al., 2015; Viena et al., 2018). In contrast, we found that suppressing PL➔PER had the largest effect on lags = −3 and the smallest effect on lags = −1. This pattern resembles a reduction in a temporal context memory retrieval strategy. Although temporal context memory has been primarily attributed to the hippocampus (Howard and Kahana, 2002; Hsieh et al., 2014; Roberts et al., 2014; Kragel et al., 2015), PER may also be involved in this memory process. For example, PER is essential in trace conditioning (Kholodar-Smith et al., 2008b; Bang and Brown, 2009) and for unitizing representations of discontinuous events (Kholodar-Smith et al., 2008a; Suter et al., 2013). Altogether these results demonstrate, for the first time, specific roles for RE and PER in memory for sequences of events, that rats use multiple retrieval strategies during sequence memory, and these strategies can be controlled by reducing the activity states of the PL➔RE and PL➔PER pathways.

Interestingly, rats showed a similar level of performance deficits following suppression of both PL➔RE and PL➔PER pathways on items that skipped ahead in the sequence (e.g., AB<u>D</u>). Theoretically, only temporal context memory or ordinal representational strategies (which were not tested here; see Orlov et al., 2000; Reeders et al., 2018) can support accurate performance when predicting upcoming items. Thus, these results show that both PL➔RE and PL➔PER pathways are critical for making predictions of upcoming items. But the exact nature of each pathway’s contribution needs to be explored in future experiments focused on items skipping ahead in sequences and/or by manipulating positional strategies.

## Conclusions

We presented evidence that top down PL pathways targeting RE and PER can control the retrieval strategy used to support sequence memory. Generally, the ability to shift memory retrieval strategies is important for situation-specific memory access and optimal memory-guided behavior. Importantly, the RE and PER pathways endow PL with the ability to exert top down control over episodic memory retrieval. In future studies, it will be worth exploring if these pathways are vulnerable in disorders that affect the temporal organization of memory such as in schizophrenia and Alzheimer’s disease.

## Acknowledgements

This work was supported by NIH grant R01 MH113626. A special thanks to all the members of the AllenLab, specifically Dr. Leila M. Allen, Dr. Tatiana Viena, and our undergraduate research assistants, John Perez, Randee Viena, and Chiara Pavon, who helped with the data collection and Amanda Rojas for her assistance with graphical illustrations of schematic renderings. We thank Dr. Aaron Mattfeld and Amanda Renfro, M.S., for useful feedback on the manuscript. Also we would like to thank the Animal Care Facility and Dr. Horatiu Vinerean DVM, DACLAM.

## Author Contributions

Conceptualization, T.A.A. and M.J.; Methodology, T.A.A. and M.J.; Investigation, M.J., M.S., S.L.; Writing – Original Draft, M.J., S.L., T.A.A.; Writing – Review and Editing, T.A.A., S.L., R.P.V., S.V.M., M.S., and M.J.; Funding Acquisition, T.A.A. and R.P.V.; Resources, T.A.A., R.P.V.; Supervision, T.A.A.

## STAR METHODS

### KEY RESOURCES TABLE

| REAGENT or RESOURCE | SOURCE | IDENTIFIER |
| --- | --- | --- |
| **Antibodies** |  |  |
| Rabbit anti-Red Fluorescent Protein | Rockland, Inc | 600-401-379 |
| Mouse Anti-NeuN Antibody, clone A60 | Millipore, Inc | MAB377 |
| VectaFluor Duet Immunofluorescence Double Labeling Kit, DyLight 594 Anti-Rabbit (red)/DyLight 488 Anti-Mouse (green) | Vector Laboratories | DK-8828 |
| Biotinylated Goat Anti-Rabbit IgG Antibody | Vector Laboratories | BA-1000 |
| VECTASTAIN Elite ABC HRP Kit | Vector Laboratories | PK-6200 |
| VECTASHIELD Antifade Mounting Medium | Vector Laboratories | H-1200 |
| VECTASHIELD HardSet Antifade Mounting Medium | Vector Laboratories | H-1400 |
| 3,3'-Diaminobenzidine | Sigma-Aldrich | CAS: 91-95-2 |
| **Bacterial and Virus Strains** |  |  |
| CAG.mCherry-2a-hM4D<sub>i</sub><sup>nrxn</sup> | AddGene | 52523 |
| AAV9.CAG.mCherry-2a-hM4D<sub>i</sub><sup>nrxn</sup>.WPRE.SV40 | UPenn Vector Core | Custom |
| AAV9.CB7.Cl.mCherry.WPRE.rBG | UPenn Vector Core | 105544-AAV9 |
| **Chemicals, Peptides, and Recombinant Proteins** |  |  |
| 1-Octanol | Sigma-Aldrich | CAS: 111-87-5 |
| (-) - Limonene | Sigma-Aldrich | CAS: 5989-54-8 |
| l-Menthone | Sigma-Aldrich | CAS: 14073-97-3 |
| Isobutyl Alcohol | Sigma-Aldrich | CAS: 78-83-1 |
| Acetophenone | Sigma-Aldrich | CAS: 98-86-2 |
| (1S) - (-) - beta-Pinene | Sigma-Aldrich | CAS: 18172-67-3 |
| L (-) - Carvone | Sigma-Aldrich | CAS: 6485-40-1 |
| 5-Methyl-2-Hexanone | Sigma-Aldrich | CAS: 110-12-3 |
| Clozapine-N-Oxide | Caymen Chemicals | ab141704 |
| Clozapine | Abcam Chemicals | ab120019 |
| **Experimental Models: Organisms/Strains** |  |  |
| Rat: Long Evans | Charles River Laboratories |  |
| **Software and Algorithms** |  |  |
| MATLAB (Version R2016a) | https://www.mathworks.com/products/matlab.html | RRID: SCR_001622 |
| OmniPlex (Version 1.17)<br>CinePlex (Version 3.6.0) | https://plexon.com/ | RRID:SCR_014803 |
| SPSS (Version 20) | https://www.ibm.com/ | RRID:SCR_002865 |
| Excel 2016 | Microsoft Office |  |
| Fluoview FV10-ASW Version 04.02.04.05 | https://www.olympus-lifescience.com/ | RRID:SCR_014215 |
| Image J 1.49v | https://imagej.nih.gov/ij/ | RRID:SCR_003070 |

## CONTACT FOR REAGENT AND RESOURCE SHARING

Further information and requests for resources should be directed to and will be fulfilled by the Lead Contact, Timothy A. Allen.

## EXPERIMENTAL MODEL AND SUBJECT DETAILS

All animal experimental procedures were conducted in accordance with the Florida International University Institutional Animal Care and Use Committee (FIU-IACUC). Subjects were male Long-Evans rats (n=34; Charles River Laboratories; weighing 250-350g upon arrival). Rats were individually housed and maintained on a 12 hr inverse light/dark cycle (lights off at 10AM). Rats had ad libitum access to food, but access to water was limited to 2–5 min each day, depending on how much water they received as reward during behavioral training (6– 9 ml). All training and testing sessions were conducted during the dark phase (active period) of the light cycle.

## METHOD DETAILS

### Sequence memory task

The sequence memory task (Allen et al., 2014) involves repeated presentations of odor-sequences and requires rats to determine whether each item (odor) was presented in-sequence (InSeq; by holding the nosepoke response for 1 sec) or out-of-sequence (OutSeq; by withdrawing their nose from the port before 1s). Rats were trained on two sequences, each composed of four distinct odors (*e.g*., Seq1: A_1_B_1_C_1_D_1_, Seq2: A_2_B_2_C_2_D_2_). Each sequence was presented at either end of linear track maze. Odor presentations were initiated by a nose-poke, and each trial was terminated after the rat either held for >1.0s (signaled by a tone) or withdrew before 1.0s (signaled by a buzzer). There was a 1.0 s interval between the start of the next trial. Water rewards (one packet of Aspartame for every 500mL of water) were delivered below the odor port after correct responses. Following an incorrect response, a buzzer sound was emitted and the sequence was terminated. Each sequence was presented 50-100 times per session in which approximately half the presentations included all items InSeq (ABCD) and half included one item OutSeq (*e.g.* AB<u>A</u>D, odor A repeated in the 3^rd^ position). Note that OutSeq items could be presented in any sequence position except the first position (*i.e.*, sequences always began with an InSeq item). Sequence memory was probed with OutSeq trials (*e.g.*, AB<u>A</u>D; one OutSeq trial presented per sequence randomly) and lag distances were analyzed to reveal the temporal order memory performance.

### Task apparatus

Rats were tested in a noise-attenuated experimental room. The behavioral apparatus consisted of a linear track (length, 183 cm; width, 10 cm; height; 43 cm) with walls angled outward at 15° and nose ports at each end, each capable of repeated deliveries of multiple distinct odors. Photobeam sensors were used to detect nose port entries. Each nose port connected to an odor delivery system (Med Associates). Odor deliveries were initiated by a nose poke entry and terminated either when the rat withdrew before 1 sec, or after 1 sec had elapsed. Water ports were positioned under each nose port for reward delivery. Timing boards (Plexon) and digital input/output devices (National Instruments) were used to measure all event times and control the hardware. All aspects of the task were automated using custom MATLAB scripts (MathWorks R2016a). A 256-channel Omniplex D with video tracking and Cineplex behavior software (Plexon) was used to interface with the hardware in real time and record behavioral data. Odors consisted of organic odorants contained in glass jars (A1: 1-Octanol, CAS: 111-87-5; B1: (-) - Limonene, CAS: 5989-54-8; C1: I-Menthone, CAS: 14073-97-3; D1: Isobutyl Alcohol, CAS: 78-83-1; A2: Acetophenone, CAS: 98-86-2; B2: (1S) - (-) - beta-Pinene, CAS: 18172-67-3; C2: L (-) - Carvone, CAS: 6485-40-1; D2: 5-Methyl-2-Hexanone, CAS: 110-12-3) that were volatilized with nitrogen air (flow rate, 2 L/min) and diluted with ultrapure air (flow rate, 1 L/min). To prevent cross-contamination, separate Teflon tubing lines were used for each odor. These lines converged into a single channel at the bottom of the odor port. In addition, a vacuum located at the top of the odor port provided constant negative pressure to quickly evacuate odor traces with a matched flow rate.

### Sequence memory task training

Naïve rats were initially trained in a series of incremental stages over 15-20 weeks. Each rat was trained to poke and hold their nose in an odor port for a water reward. The minimum required nose poke duration started at 50ms and was gradually increased (in 15ms steps) until rats held reliably for 1.2s ≥80% of the time over three sessions (200-300 nose pokes per session). Rats were then habituated to odor presentations in the port (odor A_1_ and A_2_, then odor sequences A_1_B_1_ and A_2_B_2_) and were required to maintain their nose poke response for 1.0 s to receive a reward. Next, rats were trained to identify InSeq and OutSeq items. Rats were initially trained on a two-item sequence in which they were presented with “AB” and “AA” sequences in equal proportions. The correct response on the first odor was to hold for 1.0s (Odor A was always the first item). The second response required rats to determine whether the second item was InSeq (AB; hold for 1.0s to receive reward) or OutSeq (AA; withdraw before 1.0s to receive a reward). After reaching criterion on the two-item sequence, the number of items per sequence was increased to three and four in successive stages (criterion: 70% correct across all individual odor presentations over three sessions). After reaching criterion performance on the two four-item sequences (70% correct on both InSeq and OutSeq items), rats underwent surgery for cannula implantation.

### Cannula system

A cannula implant system was created using a high-resolution (56μm) stereolithography 3D printer (ProJet 1200; 3D Systems), suitable for chronic headstages. A custom-designed 3D-printed cannula assembly was created using CAD software (Autodesk Inventor Pro Edition) and assembled with guide cannula (27 gauge, (ga).; outer diameter, (o.d.), 0.41 mm; inner diameter, (i.d.), 0.31 mm; Component Supply Company, FL) targeting PER bilaterally (A/P −6.0 mm, M/L ±6.8 mm, D/V −6.0 mm) and a single site aimed at RE (at a 10° angle in order to avoid the superior sagittal sinus; A/P −1.8 mm, M/L −1.2 mm, D/V −6.7 mm).

### AAV9 microsyringe infusions

Rats were anesthetized with isoflurane (induction 5%; maintenance: 2-3%) mixed with oxygen (800 ml/min) and placed in a stereotaxic apparatus (David Kopf Instruments, Model 900). A protective ophthalmic ointment (Gentak, 0.3%) was applied to their eyes and the scalp was sterilized with applications of isopropyl alcohol (70% in diH_2_O) followed by Betadine. The incision site was locally anesthetized with Marcaine^®^ (7.5 mg/ml, 0.5 ml, s.c.) and the skull was exposed following a fish eye incision. Adjustments were made to ensure bregma and lambda were level (±0.05μm in the D/V plane). Body temperature (35.9-37.5ºC) was monitored and maintained throughout surgery using a rectal thermometer and water heating pad. Ringer’s solution with 5% dextrose was administered to maintain hydration (5 ml, s.c.), as well as glycopyrrolate (0.2 mg/ml, 0.5mg/kg, s.c.) to prevent respiratory difficulties.

Burr holes were drilled bilaterally over PL (infusion site; OmniDrill35, World Precision Instruments). Infusions were performed using a 10 μl microsyringe (NanoFil; World Precision Instruments) and an infusion pump (UltraMicroPump III; World Precision Instruments). hM4Di+ rats (n=13) received injections of 0.5 μl of the custom AAV-hM4D_i_ ^nrxn^ (AAV9.CAG.mCherry-2a-hM4D_i_ ^nrxn^.WPRE.SV40; UPenn Vector Core) bilaterally into PL (A/P 3.24 mm, M/L ±0.7 mm, D/V from cortex −2.8 mm) at a flow rate of at 50 nl/min. mCherry-only rats (n=9) received 0.5 μl of the AAV without hM4Di (AAV9.CB7.CI.mCherry. WPRE.rBG; UPenn Vector Core) bilaterally in PL. Pilot experiments were used to determine the viral gestation time and viral expression. AAV9.hM4Di was injected in a group of rats (n=4) and perfused at a series of time points (t = +1 week, t = +2 weeks, t = +4 weeks, t = +8 weeks). In another set of rats (n=2), saline was injected into the left hemisphere of PL and AAV9.hM4Di was injected into the right hemisphere of PL, in order to determine mCherry fluorescence in the virus. The AAV9.CB7.CI.mCherry was injected into three rats at respective diultions of 1:2, 1:4, and 1:8 to measure the expression rate compared to the hM4Di+ group to determine the concentration to use for the control group.

Following injection of the viral vector into PL, burr holes overlying PER bilaterally (A/P −6.0 mm, M/L ±6.8 mm, D/V −6.0 mm), and RE (at a 10° angle in order to avoid the superior sagittal sinus; A/P −1.8 mm, M/L −1.2 mm, D/V −6.7 mm) were drilled into the skull. The cannula implant was inserted and secured with skull screws (1/8-inch grade 2 (CP) titanium; Allied Titatnium Inc.). The head stage was affixed to the surgical screws with dental cement (methyl, methacrylate, Patterson Dental). Dummies were inserted into the cannula poles (extending 0.5μm beyond the tip of the cannula) to protect against debris entering the cannula and prevent scar tissue from developing and blocking the inserted tip of the cannula. A protective cap was affixed atop the cannula implant protect from impacts and debris. Excess skin was sutured (black silk suture 4-0, with reverse cutting needle 19mm, 1/2 Circle; FEN suture). The skin surrounding the head stage was dressed with Neosporin®. At the end of surgery, rats were administered Flunixin (50mg/ml, 2.5 mg/kg, s.c.), a nonsteroidal anti-inflammatory analgesic. Rats were returned to a clean recovery incubator, and were monitored until they awoke. One day following surgery, dummies cannula were checked, and rats were administered a dose of Flunixin, and Neosporin^®^ was reapplied.

### Suppressing PL neurons and projections

According to protocol published by Roth (2016), Clozapine-n-oxide (CNO; Cayman Chemical Company) was dissolved in 0.5% DMSO in 0.9% Saline (1.0 mg/ml). CNO dose was selected based on evidence of both its behavioral effectiveness and ability to inactivate terminal activity when intracranially infused over hM4Di expressing terminals (Smith et al., 2016). The vehicle was a solution of 0.5% DMSO in 0.9% Saline. For i.p. injections, either CNO or vehicle was administered. 30-min post-injection behavioral testing commenced (randomized). For intracranial infusions, CNO and vehicle were administered over 10 min (at a volume of 1μL) in both RE and PER. Infusion cannula (32 ga.; o.d.,.0095 mm; i.d.,.005 mm; Component Supply Company) were left in place for an additional 5 minutes to allow the drug to diffuse. Behavioral testing commenced 30 minutes post infusion. Both vehicle and CNO infusions for RE and PER were counterbalanced and randomized.

### Histology

Rats were anesthetized with isoflurane (5%) mixed with oxygen (800 ml./min) and were transcardially perfused with 100 ml phosphate-buffered saline (PBS), followed by 200 ml of 4% paraformaldehyde (PFA, pH 7.4; Sigma-Aldrich). Brains were post-fixed overnight in 4% PFA and were then placed in a 30% sucrose solution for cryoprotection. Frozen brains were cut on a sliding microtome (40 μm; coronal) into 3 sets of immediately-adjacent sections. In order to visualize AAV9.hM4Diexpression as well as cannula tracts, half of the first set of slices (set 1) were mounted and cover slipped using Vectashield^®^ antifade mounting medium with DAPI. An mCherry reporter molecule was expressed with hM4Di and visualized with a confocal microscope (Olympus FV1200) using standard filter cubes. The other half of set 1 were mounted for a cell body-specific Cresyl Violet stain and coverslipped with Permount to visualize the cannula placement. A separate set of tissue was processed for immunohistochemistry to visualize the extent of PL infected terminals in each of the cannula target sites (RE and PER). Free floating sections were placed in the primary antibody: polyclonal rabbit anti red fluorescent protein (Rockland Inc.) in 0.1% bovine serum albumin (BVA) at a 1:1000 concentration for 24-48 hours. Following this, sections were washed with 0.1M phosphate buffer, placed in the secondary antibody (Biotinylated goat anti-rabbit, Vector Laboratories) at 1:500 concentrations in diluent for 2 hours. After washes, sections were incubated for 1 hour using the avidin biotin complex (ABC) Elite kit (Vector Labs) in the diluent at a 1:300 concentration. Following final washes, brown cell bodies and fibers expressing mCherry were visualized by incubating tissue in 0.022% 3,3’-diaminobenzidine and 0.003% hydrogen peroxide for approximately 2-4 min. Sections were then mounted on chrom-alum gelled slides, dehydrated in graded methanols, and placed in xylene before being cover slipped with Permount.

To map the spread of the injection, coronal micrographs were taken tissue processed for the mCherry antisera at 100x of whole slices across the frontal cortex using a NikonFI-3 mounted on a Nikon Eclipse E600 microscope for each rat (Figure S4). Micrographs were transposed over rendered plates drawn from Nissl sections and modified schematic plates (Swanson, 2004) in Procreate (Savage Software Group). These images were imported into Adobe Illustrator (Adobe Inc.) where labeled cell bodies were identified and used to pixelate and create a shaded image of the injection spread, such that intensity of shading (pixilation) corresponded to density of cell expression. The opacity of each shading was reduced to 40% and all rats were superimposed onto one another to create a final schematic at five anterior posterior levels across PL.

Lastly, a series of tissue was processed in a subset of animals (n=3) for dual immunofluorescence using antibodies for mCherry and NeuN in order to quantify the DREADD infection rate in hM4Di+ rats. For this, free floating sections were incubated in the primary antibody rabbit anti red fluorescent protein for 24-48 hours. Following PB washes, tissue was placed in an antisera for NeuN, (Mouse Anti-NeuN Antibody, clone A60; EMD Millipore) for 4 hours. Following another set of PB washes, tissue was incubated in fluorophore conjugated secondary antibodies (VectaFluor Duet Immunofluorescence Double Labeling Kit, DyLight 594 Anti-Rabbit/DyLight 488 Anti-Mouse) for an additional two hours before being mounted onto chrom-alum gelled slides, rinsed in diH_2_O, and coverslipped using VECTASHIELD HardSet Antifade Mounting Medium. Identical images of the frontal cortex were captured at 100x using epifluorescence with a NikonFI-3 camera using using NIS Elements software (Nikon) using filters for red fluorescent protein (hM4Di/ mCherry labeled neurons) and green fluorescent protein (NeuN labeled neurons) at three separate anterior posterior levels: at the core of the injection (∼3.2 from Bregma) and one level anterior and posterior (± 120um in distance). Images were then imported into FIJI Image J (Version 2.0.0; NIH), and a uniform region of interest (ROI) at each level of PL and anterior cingulate cortex, restricted to layers 5/6, which expressed the greatest density of hM4Di neurons, was selected to estimate the maximum percentage of cells infected. Manual cell counts were conducted using the Cell Counter plug in in FIJI and the ratio of mCherry labeled neurons to neurons expressing NeuN was calculated and averaged across rats.

## QUANTIFICATION AND STATISTICAL ANALYSIS

All data was analyzed in MATLAB 2016a (Mathworks), SPSS 20.0.0, and Excel 2016 using custom scripts and functions.

### Statistics

Performance on the task can be analyzed using a number of measures (Allen et al., 2014). The first position of each sequence was excluded from all analysis as these items are always InSeq. Expected vs. observed frequencies were analyzed with G-tests to determine whether the observed frequency of InSeq and OutSeq responses for a given session was significantly different than the frequency expected by chance. G-tests provide a measure of performance that controls for response bias and is a robust alternative to the χ^2^ test, especially for datasets that include cells with smaller frequencies (Sokal and Rohlf, 1995). To compare performance across sessions or animals, a sequence memory index was calculated (SMI; Allen et al., 2014) as shown in the following equation:

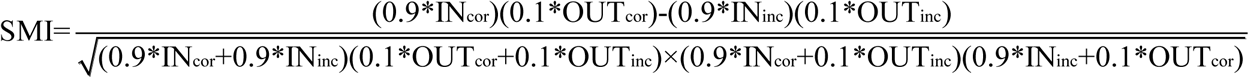

The parameters of the equation are as follows: INcor = InSeq correct, INinc = InSeq incorrect, OUTcor = OutSeq correct, and OUTinc = OutSeq incorrect. The SMI normalizes the proportion of InSeq and OutSeq items presented during a session and reduces sequence memory performance to a single value ranging from −1 to 1. A score of 1 represents a perfect sequence memory in which a rat would have correctly held their nose poke response to all InSeq items and correctly withdrew on all OutSeq items. A score of 0 indicates chance performance. Negative SMI scores represent performance levels below that expected by chance. A behavioral curve was analyzed for each rat to determine whether they performed above chance levels by measuring the SMI during each session excluding CNO infusion days. SMI was calculated to determine the effects between CNO and vehicle i.p., and infusions (RE and PER). Using SPSS, a general linear model was run to analyze the overall effects of vehicle versus CNO between hM4Di+ and mCherry-only groups. In addition, a simple linear regression was run to analyze the relationship between repeated conditions and infusions. Nose poke duration was analyzed using paired t-tests to determine whether rats held their responses significantly longer on InSeq_correct_ than OutSeq_correct_ trials. Pairwise t-tests were performed to determine any effects on inter-odor-intervals and inter-sequence-intervals between vehicle and CNO. General poke distributions were created through MATLAB using the session data.

Two distinct types of OutSeq probe trials were used to determine sequence memory: backward lags and forward lags. Backward lags occur when an odor was repeated in the sequence (e.g. AB<u>B</u>D). In this task there were three backward lags (-3-Back, −2-Back, and −1-Back). Forward lags occur when the sequence skips ahead (e.g. A<u>D</u>CD). In this particular task, there are two forward lags (+1-Fwd, and +2-Fwd). The lag analyses were used to measure performance on specific forward and backward lag trials for OutSeq items. The accuracy was calculated for each rat. Each individual rats’ vehicle performance was subtracted from their CNO performance for each lag and then divided by their difference of lag and vehicle accuracy. This produced a value that indicated the percent drop of lag performance based on each individual rats’ vehicle performance. A paired t-test was performed between RE InSeq and PER InSeq performance levels. A one-tailed, one-sample t-test was performed to measure differences against no change for all lag distances. A repeated measures 2×3 ANOVA with a Greenhouse-Geisser correction was performed on the backward lag trials to analyze any interaction effects between RE and PER infusions. A 2×2 ANOVA with a Greenhouse-Geisser correction was performed for the forward lags between RE and PER infusion. The Greenhouse-Geisser was used since in most cases the assumption of sphericity is violated for within-subjects analysis and the Greenhouse-Geisser correction is robust to this violation.

