## Supplemental Figures for "Prefrontal pathways provide top down control of memory for sequences of events"

**Supplemental Information**

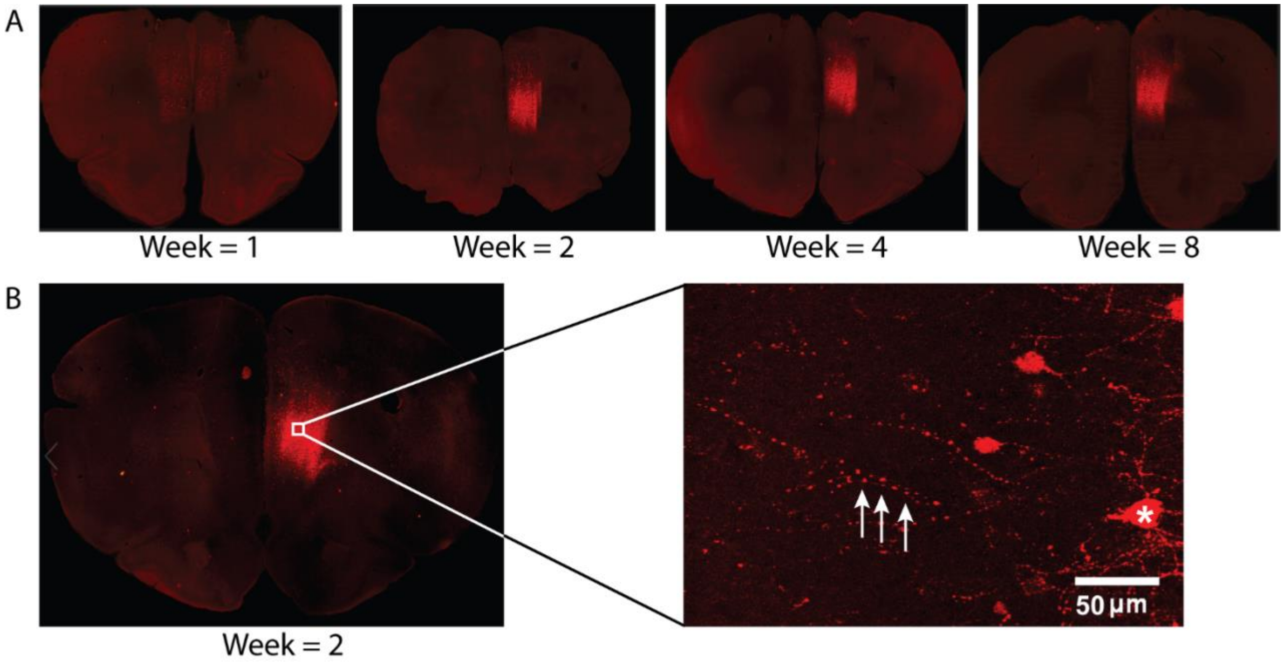


**Supplement 1. Incubation time and AAV9.hM4Di expression for these experiments.**

AAV9.hM4Di was fully expressed within two weeks within the axons and soma of neurons

(A) AAV9.hM4Di was injected unilaterally into PL in 4 rats. Each rat was perfused at different time points (weeks). The images show that by the 2-week mark, the virus was well expressed, and thus behavioral experiments began after this incubation time.

(B) AAV9.hM4Di expressed along the axons of PL neurons, as expected. The zoomed fluorescent image shows a sample axon and soma expression hM4Di, indicated by the arrows and the star respectively.

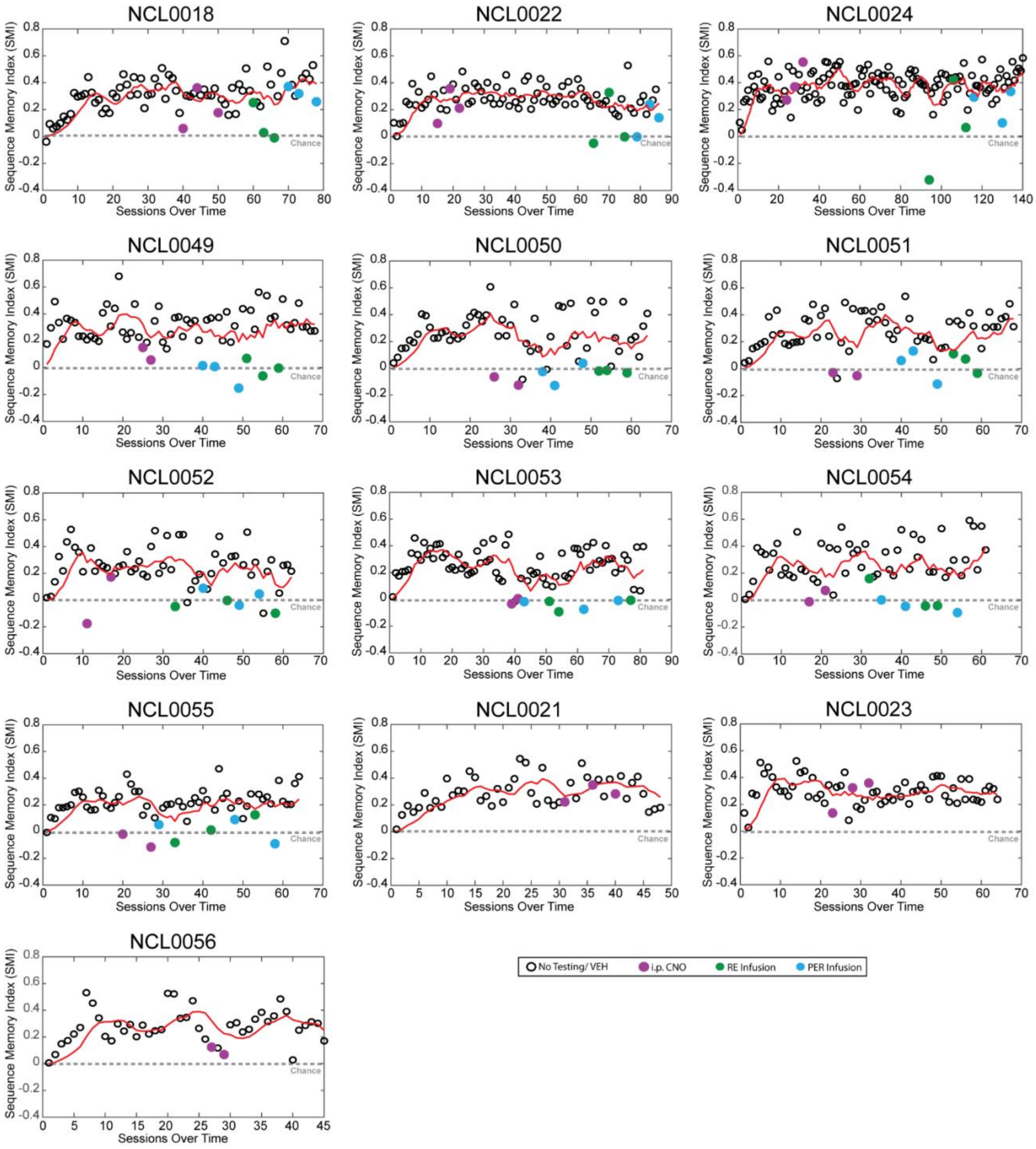


**Supplement 2. Individual behavioral performance.**

Each rat's behavioral data was collapsed into a single normalized measure (SMI) for each session. The identification of each rat is indicated at the top of each graph (NCLXXXX). The red line represents the mean SMI as the rat learned the task. All rats reached steady state and were able to maintain their SMI except for when infusions/injections occurred. With each infusion/injection there was a definite drop (at chance levels) in SMI.

Abbreviations: CNO, clozapine N-oxide; VEH, vehicle; SMI, sequence memory index; RE, nucleus reuniens of the thalamus; PER, perirhinal cortex; i.p., intraperitoneal; * = $p < 0.05$; ** $p < 0.01$; *** = $p < 0.001$.

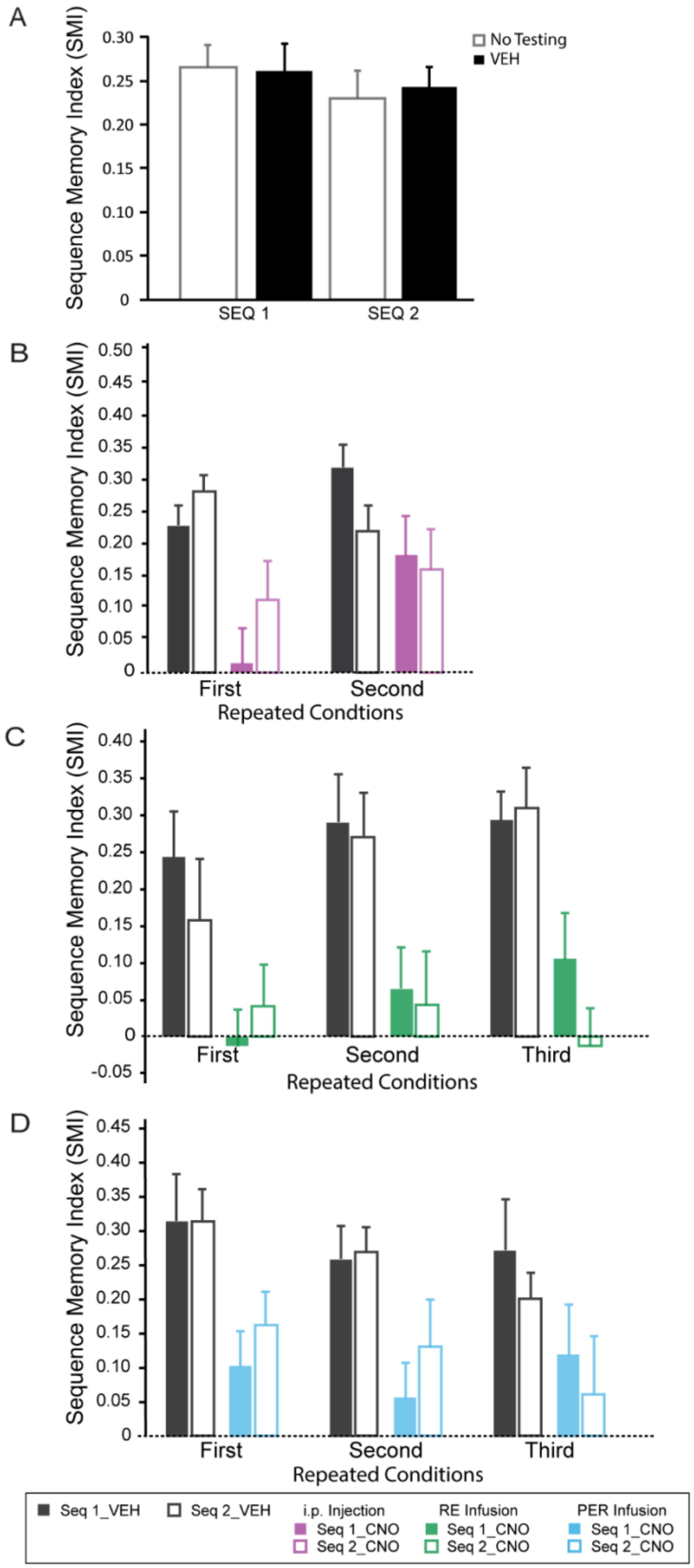


**Supplement 3. Sequence1 vs. sequence2.**
Comparison of sequence1 vs. sequence2 revealed that the rats performed at comparable levels on both sequences.
(A) Comparing normal (no testing days) vs vehicle days in sequence1 and in sequence2 indicated nonsignificant differences. Moreover, there were no significant difference between sequence1 and sequence2 for no testing days and vehicle days.
(B - D) Rats did not have a preference for sequence1 vs sequence2 during injections/infusions.
(B) There was no significant difference between sequences across repeated conditions for i.p. CNO administration.
(C) There was no significant difference between sequences across repeated conditions for PL→RE infusions.
(D) Similarly, there was no significant difference between the sequences across repeated conditions for PL→PER infusions.
Abbreviations: CNO, clozapine N-oxide; VEH, vehicle; SMI, sequence memory index; Seq1, sequence 1; Seq2, sequence 2; RE, nucleus reuniens of the thalamus; PER, perirhinal cortex; i.p., intraperitoneal; * = $p < 0.05$; ** $p < 0.01$; *** = $p < 0.001$.

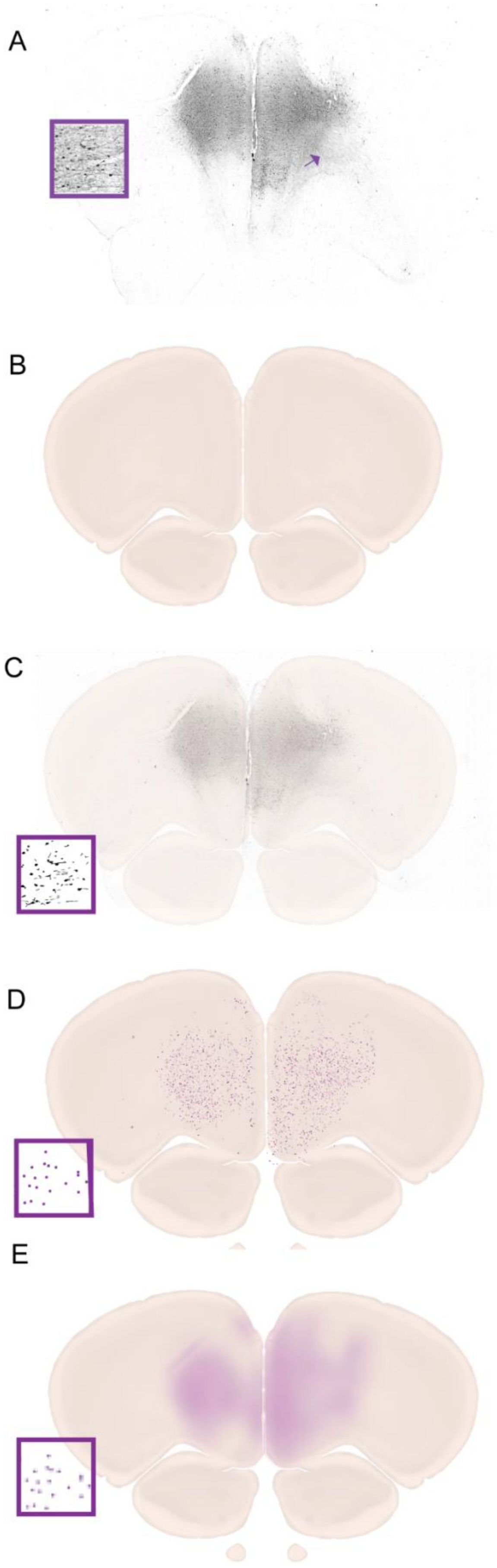


**Supplement 4. Schematic of viral spread in h4MDi+ rats**.

To illustrate how we analyzed the spread of the viral injection, a representative sample for a schematic rendering of the viral spread at one level for one rat is outlined below.

(A) Coronal micrographs were taken at 100x of whole slices across the frontal cortex from tissue processed with immunostaining (antisera for mCherry) for each h4MDi+ rat.

(B) Next, micrographs were transposed over rendered plates drawn from Nissl sections and modified schematic plates (Swanson, 2004) in Procreate (Savage Software Group).

(C) These images were imported into Adobe Illustrator (Adobe Inc.) and using the 'Image Trace' function, only labeled cell bodies were identified and used to pixelate the image

(D) This produced a shaded image of the injection spread, such that intensity of shading (pixilation) corresponded to density of cell expression.

(E) The opacity of each shading was reduced to 40% and all rats were superimposed onto one another to create a final schematic for five anterior posterior levels (see Figure 3 and S5).

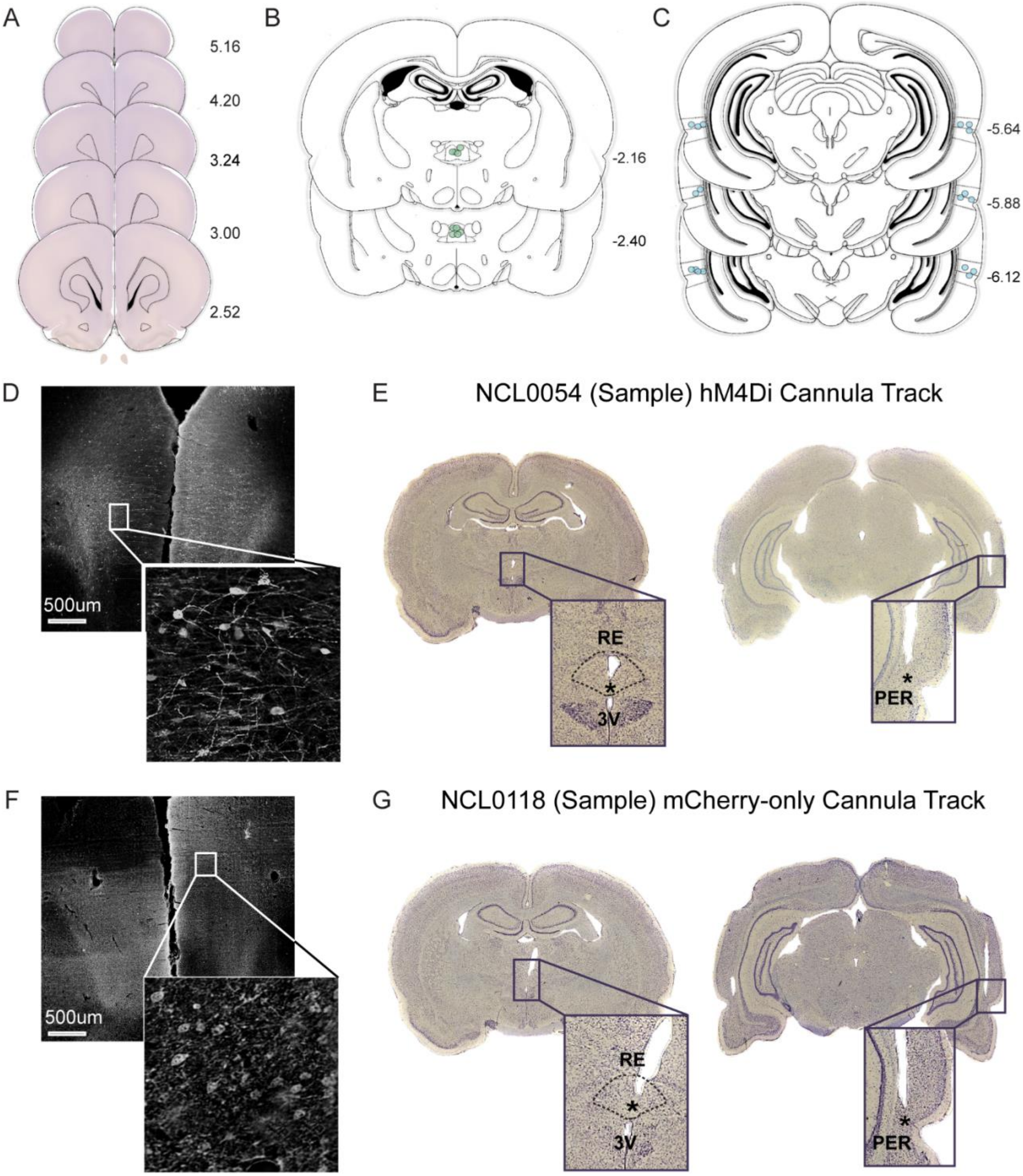


**Supplement 5. Viral expression and cannula placements for experimental groups.**

Bilateral AAV9.hM4Di or AAV9.mCherry injections were made into the PL, and guide cannula were implanted above the RE and PER (bilateral). CNO infusions targeting PL terminals in the RE or PER, effectively silencing synaptic transmission.

(A) Schematic representation AAV9.mCherry maximal viral spread in PL for all rats (n=9). While infected cells in mCherry-only rats were visualized outside of the medial wall of the prefrontal cortex, extending to the orbital and motor cortices, the null behavioral effects of these rats allowed us to confirm our findings were not associated with nonspecific effects related to the viral construct or CNO. Numbers to the right of each section indicates distance (mm) anterior to bregma according to Paxinos and Watson (2004).

(B) Microinfusion injector tip location in the RE for all AAV9.mCherry rats (n=9). Numbers to the right of each section indicates distance (mm) anterior to bregma according to Paxinos and Watson (2004).

(C) Microinfusion injector tip location for all AAV9.mCherry rats in the PER (n=9). Number to the right of each section indicates distance (mm) anterior to bregma according to Paxinos and Watson (2004).

(D) Representative image of AAV9.$hM4D_i^{nrxn}$ expression in the PL.

(E) Representative Nissl image of RE and PER (bilateral) for hM4di+. Star indicates infusion cannula tip localization.

(F) AAV9.mCherry expression in PL from a representative rat.

(G) Representative Nissl image of RE and PER (bilateral) for mCherry-only. Star indicates infusion cannula tip localization.

Abbreviations: PL, prelimbic cortex; RE, nucleus reuniens of the thalamus; PER, perirhinal cortex; CNO, clozapine-n-oxide; 3V, third ventricle.
